# Prenatal Polysubstance Exposure Alters Behaviour in Zebrafish Larvae

**DOI:** 10.1101/2025.09.19.677426

**Authors:** Lise Hermant, Gabriel D. Bossé

**Affiliations:** CERVO Brain Research Centre, 2301 Av. D’Estimauville, Québec City, QC, Canada, G1E 1T2; Department of Psychiatry and Neurosciences, Faculty of Medicine, Université Laval, Québec City, QC, Canada

**Author notes:** Correspondence: Gabriel Bossé.

**Keywords:** Zebrafish, development, sensorimotor behaviour, prenatal drug exposure, drug of abuse

## Abstract

Substance use during pregnancy has been linked to various adverse outcomes in infants, including congenital disabilities, neurodevelopmental delays, and long-term effects such as learning difficulties. An additional concern is that newborns are often exposed to multiple substances in utero. The biological consequences of such exposure remain largely unknown. Zebrafish offer an exciting alternative to fill this gap and deepen our understanding of the biological impact of prenatal multidrug exposure. We utilized zebrafish’s scalability to expose embryos to some of the most commonly used substances: nicotine, alcohol, opioids, and all their possible combinations. After embryonic drug exposure, we conducted a detailed behavioural analysis across three developmental stages. Our results revealed drug-specific outcomes, including both synergistic and antagonistic effects. Furthermore, we identified distinctive effects across development, highlighting potential developmental shifts and individual differences in resilience. Overall, these findings demonstrate that prenatal polydrug exposure results in complex, stage-dependent effects, sometimes antagonistic, which cannot be predicted from single-drug outcomes. Our study emphasizes the value of zebrafish as a model for investigating polydrug interactions and provides a framework for exploring biomarkers of vulnerability and resilience in offspring.

## 1. Introduction

Substance use during pregnancy is becoming a significant public health issue, with increasing rates of opioid, tobacco, and alcohol consumption over the past few years, and often continuing throughout pregnancy(CDC 2025). Between 2008 and 2018, the number of newborns diagnosed with Neonatal Opioid Withdrawal Syndrome increased by 242%(Conradt et al. 2018). Additionally, it is estimated that nearly 20% of pregnancies involve alcohol consumption during the first trimester alone(England et al. 2020). Prenatal drug exposure poses significant risks for maternal and infant health.

It is linked to a broad spectrum of health risks, including preterm birth(Baer et al. 2019), low birth weight(CDC 2025), sudden infant death syndrome(Makarious et al. 2022), and neonatal abstinence syndrome(Barry et al. 2021). Remarkably, 75% of drug-exposed infants experience major medical issues, compared to 27% of their unexposed peers(Huestis and Choo 2002). Beyond these early complications, children exposed to drugs in utero often face long-lasting challenges such as cognitive impairments, developmental delays, congenital abnormalities, metabolic disruptions, and an increased risk of substance use disorders later in life(Ross et al. 2015; Harder and Murphy 2019; Abu and Roy 2021; Derme et al. 2024). At the molecular level, prenatal exposure disrupts critical developmental pathways, affecting systems involved in the regulation of circadian rhythms(Pačesová et al. 2021), neuronal development(Stankovic and Colak 2022), and cardiac function(Janardhan et al. 2022; Weeks et al. 2024).

Compounding the risk is the frequent co-use of multiple substances. Among pregnant women who use non-medical opioids, 88.9% report using at least one additional substance(Jarlenski et al. 2017), most often cigarettes (42%) and alcohol (51.2%). Overall, about 5% of newborns in the United States are exposed to multiple substances *in utero*, thus affecting nearly 180,000 newborns each year(Qato et al. 2020). Polydrug exposure can exacerbate adverse outcomes compared to single-drug use, including increased rates of preterm birth(Garrison-Desany et al. 2020), and particularly severe effects when opioids are involved(Isaacs et al. 2021). Despite its prevalence, the biological consequences of prenatal polydrug exposure remain poorly understood both behaviourally and molecularly, as most research, whether clinical or preclinical, tends to focus on single-drug exposure.

In this context, zebrafish *(Danio rerio)* offer an innovative alternative to mammalian models in neuroscience research. They possess a fully developed central nervous system and a blood-brain barrier, and their brain structure and neurotransmitter systems, including inhibitory GABAergic and excitatory glutamatergic circuits, are well-conserved(Higashijima et al. 2004). Their external fertilization, rapid development, genetic accessibility, and suitability for high-throughput behavioural assays make them uniquely suited for studying the developmental effects of drug exposure in a scalable mode(Kalueff et al. 2016, 2017).

Zebrafish larvae are sensitive to a wide range of substances, including ethanol(Tran et al. 2017), nicotine(Wronikowska et al. 2020) and opioids(Lau et al. 2006), which elicit pharmacological responses similar to those observed in humans. Moreover, they display a rich and evolving behavioural repertoire as early as 3-5 days post-fertilization (dpf), showcasing various sensorimotor responses, habituation, fear responses(Rennekamp et al. 2016; Basnet et al. 2019; Bosse et al. 2021), and, by the third week, more complex behaviours such as social interactions and associative learning(Dreosti et al. 2009). Previous zebrafish studies have described developmental outcomes of single prenatal drug exposures, including cranial malformations, central nervous system defects, and aberrant apoptotic cell death after ethanol(Weeks et al. 2020; Alsakran and Kudoh 2021), altered social behaviour and elevated anxiety levels after opioids(Wang et al. 2022), and dose-dependent toxicity and growth restrictions after nicotine(Parker and Connaughton 2007). However, no study to date has systematically examined the behavioural consequences of combined exposure to these commonly co-used substances.

Here, we examine the behavioural effects of prenatal exposure to ethanol, nicotine, and morphine, three substances often co-used during pregnancy, either separately or together. Using zebrafish, we evaluated behavioural outcomes at multiple developmental stages to determine whether polydrug exposure results in additive, synergistic, or antagonistic effects. Our findings show drug-specific, time-dependent behavioural changes, underscoring the importance of modelling prenatal polydrug exposure and highlighting zebrafish as a valuable model for understanding its complex biological impacts.

## 2. Methods

All experiments described in this study were approved and performed in compliance with the *Comité de Protection des Animaux* (CPAUL) of Université Laval.

### 2.1 Embryo collection and development

Adult zebrafish (*Danio rerio*) of the wild-type Tüpfel long fin (WT-TL) line were maintained in the fish facility at 28−29°C with a 14/10 h light/dark cycle. Embryos were obtained from multiple mating pairs, pooled and randomly assigned to treatment groups. They were raised in E3 solution (NaCl 4.97 mM, KCl 0.179 mM, CaCl2 0.33 mM, MgCl2 0.40 mM, Hepes 20 mM) at a pH of 7.2 in 100 mm x 25 mm petri dishes (Avantor Science Central, USA) at a density of 1 embryo/ mL and kept at 28.5°C with a 14/10 hours light/dark cycle. The medium (with or without treatment) was renewed daily, and mortality was monitored on a daily basis. At 3 dpf, chorion debris were removed.

From 5 to 10 dpf larvae were fed daily with 30 μL/larvae of eukaryotic *Tetrahymena termophila. T. thermophila* cultured in an SPP medium enriched with peptone protease, and Fe3+ (2% proteose peptone, 0.1% yeast extract, 0.2% glucose, 33 μM FeCl3, 250 μg/mL Streptomycin/penicillin, 0.25μg/mL Amphotericin B) until their concentration reached a plateau. An 11 mL volume of SPP medium was inoculated with 1 mL of *T. thermophila* culture at maximum confluence. Before use, *T. thermophila* were washed in E3 medium from their SPP medium to remove antibiotics, using three centrifugations (2 min, 0.8 g).

At 10 dpf, larvae were transferred into 1.1 L recirculating tanks filled with deionized water supplemented with Instant Ocean Sea Salt (Tropic Marin, Wartenberg, Germany) and NaHCO₃ to maintain a conductivity of 700–1000 μS and a pH of 7.2–7.4. Larvae were housed in groups of 25–30 individuals per tank and maintained at 28.5°C under a 14:10 light-dark cycle. Water temperature, conductivity, and ammonia levels were monitored daily. From 10 dpf onward, larvae were fed daily with 24-hour-hatched *Artemia nauplii* until the end of the experiment. The amount of food was based on intake for the first 10 minutes to ensure it was not limited in quantity, as described in Caperaa *et al*. 2024(Caperaa et al. 2025).

### 2.2 Drug treatment

Embryos were randomly allocated to treatment groups, and drugs were diluted directly in E3 medium. Final concentrations were: Ethanol 1% (volume/volume) (Les alcools de commerce, Canada), nicotine 10 µM (Sigma-Aldrich, Germany) and morphine sulphate at 10 mg/L (Toronto Research Chemicals, Canada). Treatments were applied from 3 hours post-fertilization (hpf) until 72 hpf with daily renewal. At each medium change, viability was assessed by visual inspection. At 72 hpf, the drug solutions were replaced with fresh drug-free E3 medium. Embryos that had not yet hatched were mechanically stimulated to hatch by pipetting using a 5 mL pipette. Untreated siblings underwent identical handling and medium changes.

## 3. Behaviour assessment

All experiments were conducted between 9:00 a.m. and 5:00 p.m., except for the sleep test, which was performed overnight. Before testing, the larvae were transferred to the experimental room and incubated for 30 minutes at 28.5°C under a 14:10 light:dark cycle. Behavioural assays were performed in individual wells, with plate type specified in each assay section.

Larvae were transferred to an experimental plate, which was then loaded into a *Zebrabox* system (ViewPoint, France) equipped with a computer-controlled light box and an infrared-filtered video camera. For each session, data were acquired at 30 frames per second (fps), except for the sleep test, which was performed at 10 fps. The same larvae were tested at 8, 14, and 21 dpf with 12 animals per batch and a minimum of two independent batches. Locomotor activity was quantified using the quantization method, based on pixel difference, from the *Zebralab* software (ViewPoint, France) with the following settings: background refresh = 300, sensitivity= 20.

### 3.1.1 Multi-stimuli assay (MSA)

This protocol was designed to evaluate larval responses to a range of sensory stimuli and is described in detail in Caperaa *et al*. (2025)(Caperaa et al. 2025). Briefly, the MSA consists of a 60-minute experimental sequence during which animals are exposed to various stimuli presented in a predefined order, alternating between periods of darkness and full-spectrum white light, mechanical vibrations, and light flashes of varying intensities. At 8 dpf, larvae were tested individually in 96-well plates (Corning, USA) with 200 µL of E3 medium per well. At 14 dpf, larvae were tested in 24-well plates (Sarstedt, Germany) using 500 μL of home-tank water per well.

### 3.1.2 Light-dark preference

This test evaluated larval activity before, during and after exposure to a stressful strobe-light stimulus(Rennekamp et al. 2016; Caperaa et al. 2025). Experiments were conducted using a 6-well plate (Greiner Bio-One, Austria), with each well partially covered with black optic tape (on the lid and the bottom) to create light and dark zones, allowing larvae to remain in the illuminated area or hide in the dark. Each well contained a single larva and was filled with either 5 mL of E3 medium (8 dpf) or home-tank water (14 dpf). The experimental sequence consisted of 10 minutes of full-spectrum white light (baseline), 3 minutes of strobe light exposure (1 minute of darkness followed by 1 minute of 10Hz strobe light). It concluded with 10 minutes of full-spectrum white light to assess post-stress recovery. Larvae previously tested in the MSA were excluded from this test to avoid repeated exposure to overlapping stimuli.

### 3.1.3 Sleep test

Zebrafish possess a circadian rhythm and are highly sensitive to light, with the transition to darkness typically triggering sleep-like behaviour (Hurd *et al.,* 1998). To monitor circadian activity, larvae were recorded overnight for 18.5 hours. At 8 and 14 days post-fertilization (dpf), larvae were placed individually in 24-well plates (Sarstedt, Germany) with 2 mL of E3 medium at 8 dpf or home-tank water at 14 dpf. The experiment followed the same light/dark cycle as used in the animal facility. It consisted of three phases: 5 hours of full-spectrum white (baseline), 10 hours of darkness (nighttime rest) and 3.5 hours of light (morning recovery). Recordings were initiated around 4 p.m. each day and acquired at a rate of 10 fps.

### 3.1.4 Thermosensation assay

At 8 and 14 dpf, larvae were briefly placed on ice in a 6-well plate until they became immobile (typically after approximately 10 minutes). They were then individually transferred to custom opaque plates (dimension comparable to standard plates) containing either 200 µL E3 medium (96-well plate, 8 dpf) or 1 mL home-tank water (24-well plate, 14 dpf). Plates were then placed on a pre-heated hot plate (50°C) and larval movement was recorded continuously for 10 minutes.

### 3.1.5 Social preference test

Social behaviour was assessed at 21 dpf, when zebrafish reliably display social behaviour preference (adapted from Dreosti *et al.,* 2015)(Dreosti et al. 2015). The U-shaped arena was divided into four zones in which the tested animal could freely swim: a “social” zone in one arm of the arena, a “non-social” zone in the opposite arm, and two “neutral” zones located between them. Observation chambers were located at the end of each arm and were separated from the main arena by transparent glass windows, allowing visual but not physical interaction. The one adjacent to the social zone contained the interactors during the test session.

Individual larvae were placed in the arena and allowed to freely explore for a 15-minute habituation period. Following this habituation, two conspecific interactors (from the same tank and of comparable size) were introduced into the observation chamber adjacent to the arm least visited by the tested larva during habituation to minimize side bias. A 10-minute test session was then recorded, and a social interaction score was calculated as follows: [(time spent in social zone during test/duration of test) x 100] – [(time in social zone during habituation/duration of habituation) x 100].

### 3.2 Quantization method

Larval activity was quantified using the *Zebrabox system* (Viewpoint, France). Recordings were made at 30 fps (MSA, light-dark preference test, social preference test) or 10 fps (sleep test). The *Zebralab* software estimated movement using pixel-intensity differences between consecutive frames. All experiments were performed under full-spectrum white light unless specified otherwise. Detection settings were consistent across assays (background refresh = 300; sensitivity = 20)

### 3.3 Statistics and data analysis

Raw data generated by the *Zebrabox* were analyzed using custom R scripts. To control for variability across clutches and days, log 2 fold-change values relative to control siblings were calculated for each treated larva at each time point. Since we always include control animals in each experiment, the log2 fold change is relative to the control tested on the same day as the drug-exposed animals. We then combine the data from all the drug treatments and perform the remaining analysis on that dataset. Data were tested for normality using the Shapiro-Wilk test, and outliers were identified using Rosner’s test. Statistical analyses are indicated in each figure legend. *P-*values were corrected for multiple comparisons using a false discovery rate analysis and Benjamin-Hochberg correction, with significance set at *P*-value < 0.05.

## 4. Results

### 4.1 Modelling prenatal polydrug exposure in zebrafish

To investigate the biological impact of polydrug exposure, we have selected three of the most consumed and abused substances: ethanol (E), morphine (M) and nicotine (N) and exposed zebrafish embryos during the first 72 hpf. This period represents a critical developmental window during which major neuronal pathways begin to form, mirroring key aspects of early fetal development in humans(Higashijima et al. 2004; Veldman and Lin 2008). Although a direct equivalence is debated, some studies suggest that the first 72 hpf in zebrafish approximate at least the first trimester of human pregnancy(Nakajima et al. 2021; Raghul Kannan et al. 2024). At 72 hpf, embryos were transferred to drug-free embryo media and raised under normal conditions.

Embryos were treated with each drug individually and all possible combinations for a total of 8 experimental conditions. We selected a single concentration for each substance, which was determined based on pilot studies conducted within our group and the existing literature(Svoboda et al. 2002; Joya et al. 2014; Sales Cadena et al. 2021; Borrego-Soto and Eberhart 2022).

To evaluate the impact of such exposure, we conducted a battery of behavioural assays at three key developmental stages: 8, 14 and 21 dpf. At 8 dpf, larvae display a diverse behavioural repertoire, including sensorimotor responses and non-associative learning (habituation)(Basnet et al. 2019),(Patton et al. 2021) and these responses evolve during development. Notably, these larval stages have well-defined neuroanatomical structures and neurotransmitter systems that are highly conserved with those of humans(Patton et al. 2021). Additionally, we also performed a social preference assay at 21 dpf when zebrafish begin to robustly exhibit social behaviours(Dreosti et al. 2015; Stednitz and Washbourne 2020).

### 4.2 Drug combination increases lethality

At 3 hpf, eggs were visually inspected to remove unfertilized or abnormally developing eggs(Kimmel et al. 1995). This step was repeated daily throughout the treatment period, during which survival was also monitored to assess the potential toxicity of prenatal drug exposure. As expected, some mortality was observed across all conditions, including the control group. Survival rates following single drug exposure were comparable to those of control animals. However, the combination of ethanol-nicotine and ethanol-morphine-nicotine resulted in a marked increase in mortality within the first 24 hours, with only 30% of embryos surviving until the end of the exposure time **(Fig.1)**.

**Figure 1:**
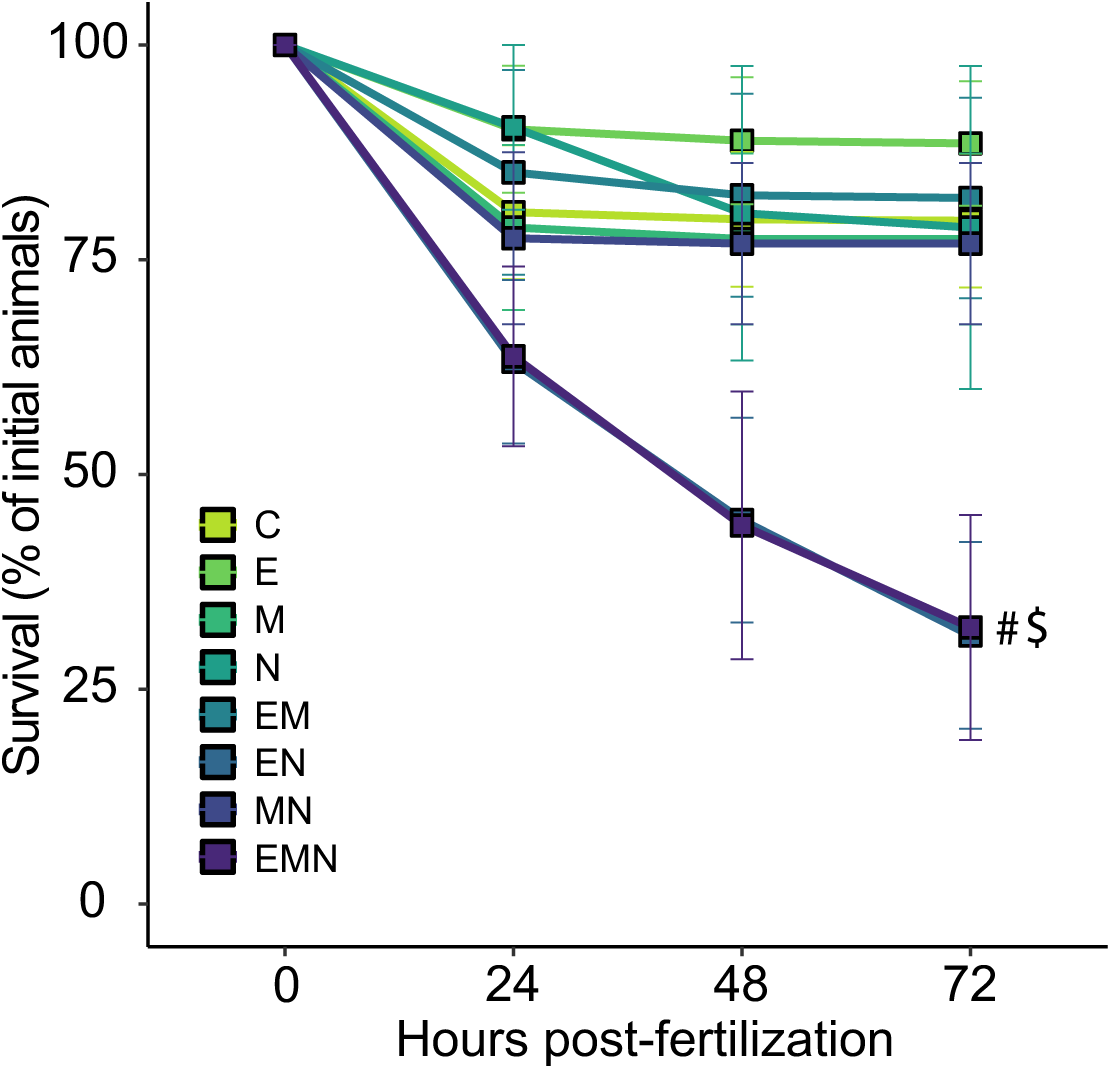
Polydrug-exposed embryos had a reduced survival rate. Survival curve for the duration of the treatment. Each dot represents the percentage of surviving embryos per condition at 24-hour intervals. Minimum of 2 batches. n >2 per condition. *P*-value computed by Tukey’s HSD significant difference test on 1-way ANOVA [F(1,100) = 221.12, *P =* 2E-16]. n = 60 per condition. #p <0.05 for the EN group when compared to controls. $p <0.05 for the EMN group when compared to controls at 72 hpf. This experiment employed a between-subjects design.

### 4.3 Prenatal drug exposure affects sensorimotor responses

To measure a diverse set of sensorimotor responses, we utilized an assay developed by our group, which consists of various stimulus patterns referred to as the multi-stimuli assay (MSA)(Caperaa et al. 2025). This assay included baseline locomotion recordings in both darkness and light conditions, followed by sequences of light and vibration stimuli designed to evoke specific behavioural responses related to locomotion, fear (strobe light), stress (dark-light transitions), and sensory sensitivity (light flash). We also evaluated non-visual sensory responses through vibrations to measure mechanical sensitivity (single vibration event) and non-associative learning (habituation to repeated stimuli). Notably, these stimuli are widely used in zebrafish behavioural studies and have been shown to consistently evoke robust and reproducible responses (**Fig.2A**), which our group and others have previously described(Burgess and Granato 2007; Irons et al. 2010; Rennekamp et al. 2016; Bosse et al. 2021; Caperaa et al. 2025). As a preliminary analysis of our data, we first compared the average activity of drug-exposed larvae across the entire duration of the MSA (**Supp.Fig.1),** revealing interesting patterns. For instance, E-, M- and EM-exposed larvae exhibited an overall hyperactive phenotype, whereas N-exposed animals showed the opposite trend. Drug-paired treated groups, on the other hand, had mixed effects. For instance, EN and EMN groups showed drastic overactivity in response to specific stimuli (**Supp.Fig.1).**

**Figure 2:**
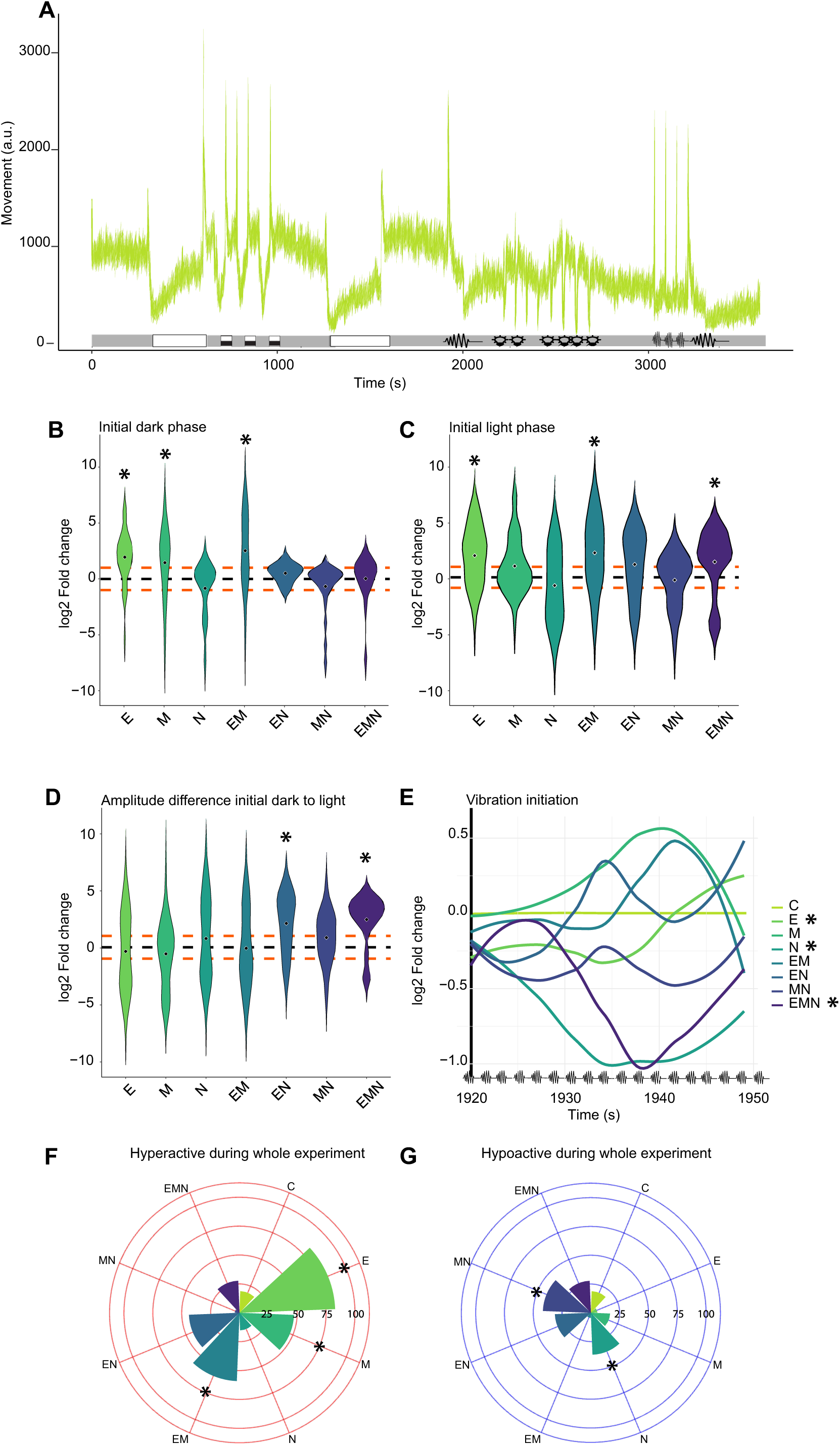
Sensorimotor responses to the Multi-Stimuli Assay (MSA) in 8 dpf larvae prenatally exposed. **A.** Average locomotor responses of control larvae (C) throughout the duration of the MSA**. B.** During the initial dark phase, larvae exposed to ethanol (E), morphine (M), and the combination of both drugs (EM) were hyperactive compared to controls. Nicotine (N)-exposed larvae showed the opposite trend, although it was not statistically significant (p = 0.18). * = p < 0.05 compared to controls. *P-*values were calculated using the Mann–Whitney test followed by Benjamini– Hochberg correction for multiple comparisons. **C**. During the initial light phase, larvae from the E, EM and EMN groups displayed hyperactive behaviour. * = p < 0.05 compared to control animals. *P-*values were calculated using the Mann–Whitney test followed by Benjamini–Hochberg correction for multiple comparisons. **D.** Transition from dark to light triggers an initial burst in movement. To quantify this, we measured the amplitude following this transition, which revealed that EN and EMN-exposed larvae responded more strongly than control larvae. * = p < 0.05 compared to controls. *P-*values were calculated using the Mann–Whitney test followed by Benjamini–Hochberg correction for multiple comparisons. **E.** Repeated vibration exposure leads to habituation and a reduction in response. To determine whether drug-exposed larvae exhibited different habituation rates, we performed a linear mixed-model analysis to assess changes in log2 fold change in response to vibration stimuli. This figure represents the average log2 fold change for each condition over time using a smoothing function. The linear mixed-model analysis revealed that E-exposed larvae showed a significant increase in log2 fold change over time, N and EMN groups displayed strong initial responses that gradually declined over successive pulses. * p < 0.05 compared to control animals. *P-*values were obtained using a linear mixed-effects model with condition over time as a fixed effect and subject (larva ID) as a random effect. **F.** The percentage of larvae with an average log2 fold change greater than 1 across the entire duration of the experiment was calculated. Most of the E- and some of the M- and EM-exposed animals were hyperactive.* = p < 0.05 compared to control animals. *P-*values were calculated by using a logical regression. **G**. The percentage of larvae with an average log2 fold change of lower than -1 across the whole duration of the experiment was calculated. N and MN-exposed were significantly hypoactive. * = p < 0.05 compared to control animals. *P-*values were calculated by using a logical regression. For all experiments, n ≥ 24 per condition. The violin plot represents the data in log2 fold change compared to control animals. The black dashed line indicates the average of control larvae (0), while the orange lines represent a log2 fold change of +1 and -1. A log2 fold change of +1 represents a twofold increase in movement, while -1 indicates a twofold decrease compared to control animals. This experiment employed a between-subjects design.

Based on these traces, we decided to quantify specific parameters, including both the initial dark phase (**Fig.2B**) and the light phase (**Fig.2C)** of the experiment. During those phases, E, M, and EM-exposed larvae exhibited a hyperactive phenotype, while N- and MN-exposed larvae tended to be hypoactive. As the traces suggested, multi-drug exposure increased reactivity to light changes (**Fig.2D)**. As part of the multi-stimuli assay, we also performed a series of rapidly repeating short bursts of vibration to assess habituation, a simple form of non-associative learning in which animals gradually reduce their response to repeated stimuli **(Supp.Fig.2A).** Control larvae responded strongly to the initial vibrations followed by a progressive decline in reactivity. To quantify the initial response, we measured the average log2 fold change immediately before the start of the vibration sequence compared with the first 3 seconds following its initiation. This revealed a blunted reaction in E- and EM-exposed larvae relative to controls, with similar but non-significant trends in EN- and EMN-exposed groups **(Supp. Fig. 2B)**. To evaluate responses across the entire stimulus sequence, we applied a linear mixed-effects model. E-exposed larvae showed a significant increase in log2 fold change over time, suggesting a sustained reactivity and impaired habituation. In contrast, N- and EMN-exposed animals displayed a more pronounced reduction in responsiveness than controls **(Fig.2E).**

To capture behavioural diversity beyond mean responses, we quantified individual-level activity patterns. For each larva, we calculated the average log2 fold change across the MSA and classified animals as hyperactive or hypoactive when their log2 fold change exceeded +1 or fell below -1, respectively (i.e., a two-fold increase or decrease relative to controls). This analysis revealed that over the duration of the MSA, 83% of E-exposed, 47% of M-exposed, and 59% of EM-exposed larvae were hyperactive (**Fig.2F),** whereas 36% of N- and 42% of MN-exposed animals were hypoactive **(Fig.2G).** During this analysis, we observed that drug-exposed animals exhibited more variable responses. We thus assessed variance within each treatment group across the whole trial. Single-drug exposures produced greater behavioural variability than controls. At the same time, this effect was attenuated in polydrug groups, suggesting that combined exposures may dampen the range of behavioural responses, leading to more uniform activity profiles **(Fig.3A)**.

**Figure 3:**
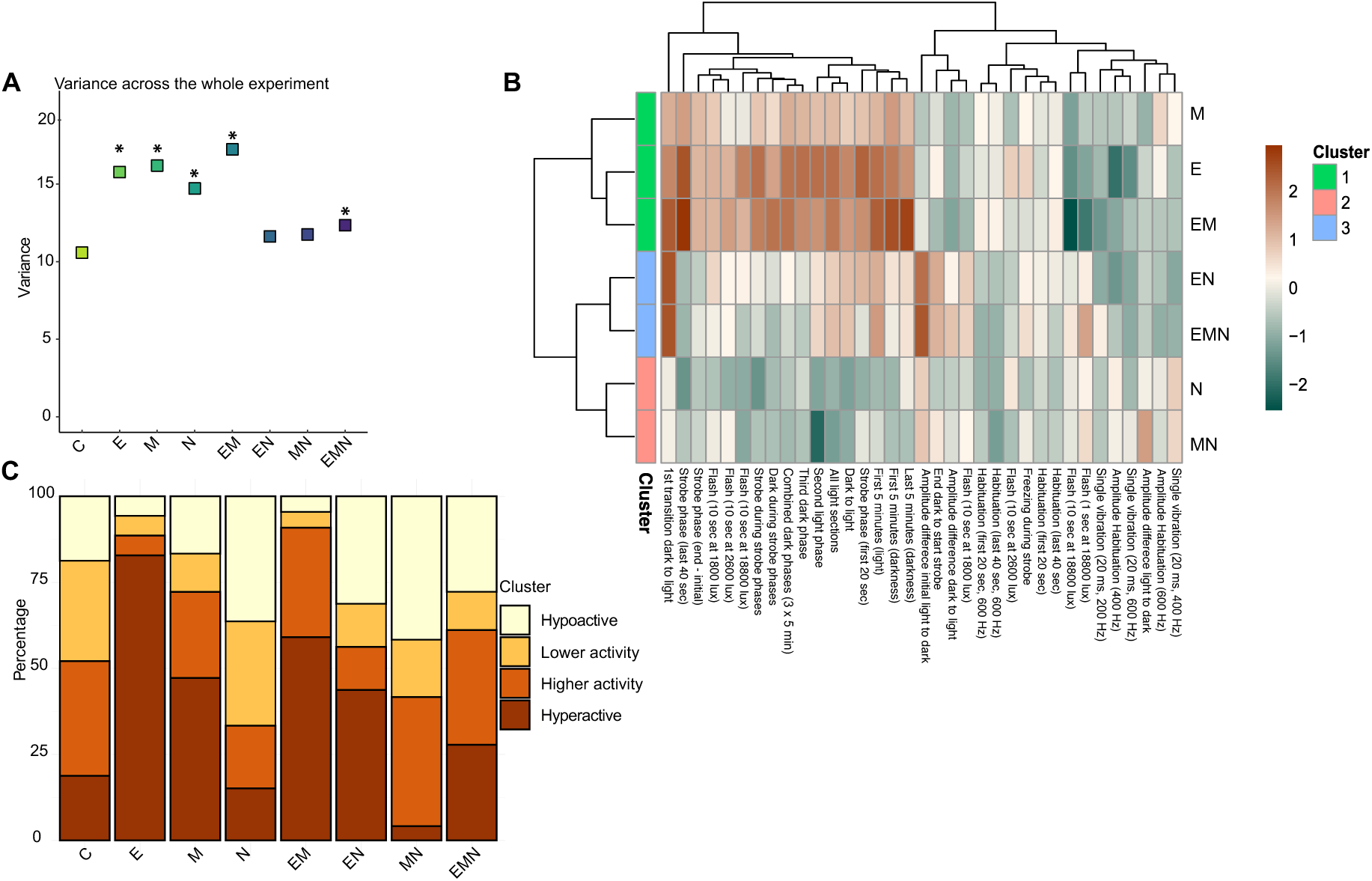
Variance and global activity in response to each stimulus of the MSA in 8 dpf larvae prenatally exposed. **A.** Animals from single drug-exposed groups and from the EM and EMN groups had a greater variance across the duration of the MSA. * = p < 0.05 compared to the control group. Significance was calculated with a Levene test followed by a Bonferroni correction for multiple comparisons. **B.** Hierarchical clustering of the different treatments (y-axis) based on the log2 fold change across the various parameters we measured. Heatmap colours indicate the log2 fold change compared to controls, where orange and blue reflect increased and decreased responses, respectively. Clusters were then generated based on similar responses. This analysis revealed three min clusters: one grouping M, E and EM animals, a second including EN and EMN animals and a third composed of N and MN animals. **C.** Manual clustering based on the average log2 fold change for each larva across the duration of the MSA. The categories were separated as follows: Hypoactive log2 fold change lower than -1, Lower activity log2 fold change between -1 and 0, Higher activity log2 fold change between 0 and 1 and Hyperactive log2 fold change greater than 1. Control animals showed a homogenous distribution, while E- and EM-exposed larvae were primarily found in the hyperactive cluster. Almost no hyperactive individuals were observed in the MN group. n ≥ 24 per condition. This experiment employed a between-subjects design.

To better understand the response distribution of drug-exposed larvae, we classified each larva based on its overall response, dividing them into four categories according to their average log2 fold change. This approach revealed different distributions across conditions. E, M and EM exposures increased the proportion of hyperactive animals, while the EN group included both hyper- and hypoactive subgroups. Interestingly, almost no MN larvae were classified as hyperactive **(Fig.3B)**. To capture broader response patterns, we also performed hierarchical clustering across all MSA parameters. This enabled the classification of each drug-exposed group into a cluster based on their responses. Notably, M, E and EM-exposed animals clustered together and showed a general trend of hyperactivity. EN and EMN-exposed larvae formed another cluster with more variable effects, while N- and MN-exposed larvae clustered together, displaying consistent hypoactive responses across most stimuli (**Fig.3C)**. Together, these findings from the MSA suggest that drug combination-specific effects shape sensory processing in exposed larvae at 8 dpf, with both enhanced variability and distinct behavioural phenotypes depending on the treatment.

To assess whether these phenotypes persisted over time, we repeated the MSA at 14 dpf using the same group of animals. A preliminary analysis revealed extremely divergent responses in a subset of MN-exposed animals (data not shown). Rosner’s test confirmed these outliers, which were excluded from further analyses. Notably, such deviations were not observed in other treatment groups from the same clutch.

At 14 dpf, average log2 fold change traces across the MSA did not reveal a consistent hyperactive phenotype in any of the drug-exposed animals. Still, EN-exposed larvae tended to display hypoactivity (**Supp.Fig.3**). As previously established, overall sensorimotor responses were comparable to 8 dpf larvae except for dark-to-light transitions, which reliably elicited a freezing reaction followed by a burst of activity (**Supp. Fig.4A**) consistent with previous findings(Caperaa et al. 2025).

Over the whole duration of the MSA, we found that 33% of E- and 38% of EM-exposed larvae displayed an average log2 fold change greater than 1. In comparison, 33% of EN-exposed larvae fell below -1, although this proportion was not significantly different from controls **(Supp. Fig. 4B-C)**. Then, phase-specific analysis revealed that E-, M- and EM-exposed larvae tended to be hyperactive during the light phase, whereas N-exposed groups, particularly EN, were more often hypoactive **(Supp.Fig.4D)**. Dark-to-light transitions triggered stronger increases in activity in the E and EM groups compared to controls **(Supp.Fig.4E)**. During the strobe light phase, E-, M- and EM-exposed larvae also displayed elevated activity **(Supp.Fig.4F).**

For the repeated vibration stimuli, initial startle responses were comparable across groups. However, linear mixed-effects modelling indicated that N-exposed larvae showed a faster decline in responsiveness over time, suggesting enhanced habituation (**Supp.Fig.4G)**. As at 8 dpf, inter-individual variance remained elevated in most drug-exposed groups except in the triple-exposed condition, which showed reduced variance (**Supp.Fig.5A**). Cluster analysis revealed broadly similar patterns to 8 dpf, with fewer hyperactive animals overall (**Supp.Fig.5B**). Finally, a hierarchical clustering of responses across all parameters grouped M-, E- and EM-exposed larvae (hyperactive) and N- and EMN-exposed larvae together (hypoactive). MN larvae were excluded from this clustering because their strong hypoactivity skewed the scale and disrupted the separation of the other groups (**Supp.Fig.5C)**.

In summary, behavioural differences were less pronounced at 14 than at 8 dpf. Still, exposure-specific trends persisted: E-, M-, and EM-exposed larvae remained predominantly hyperactive, whereas N-exposed groups tended toward hypoactivity.

### 4.5 Sleep patterns are affected in drug-exposed larvae

Alterations in circadian rhythm and sleep cycles have been reported in human newborns and adolescents following *in utero* exposure to substances such as nicotine(Stéphan-Blanchard et al. 2008) and alcohol(Chandler-Mather et al. 2021). In animal models, these disruptions have been linked to changes in the expression of key regulators of the circadian rhythm(Farnell et al. 2008; Pačesová et al. 2021).

To assess the impact of prenatal drug exposure on sleep-like behaviour, we conducted long-term movement tracking over 18.5 hours aligned with the standard light-dark cycle of our fish facility. Larvae were recorded for 5 hours during the day, followed by 10 hours of darkness (night phase), and the recording ended the next morning. At 8 dpf, control animals exhibited a gradual increase in locomotion during the day, peaking approximately 150 minutes before, and then steadily declining. Lights-off elicited a brief startle response, followed by a sustained reduction in activity punctuated by short arousals, consistent with sleep-like behaviour (**Fig.4A**). Comparing average traces across conditions revealed drug-specific differences: E, M and EM larvae displayed initial hyperactivity, while N, EN and EMN larvae showed reduced activity. During the night, M-exposed larvae were notably hyperactive, whereas N-exposed groups remained hypoactive (**Supp.Fig.6**).

**Figure 4:**
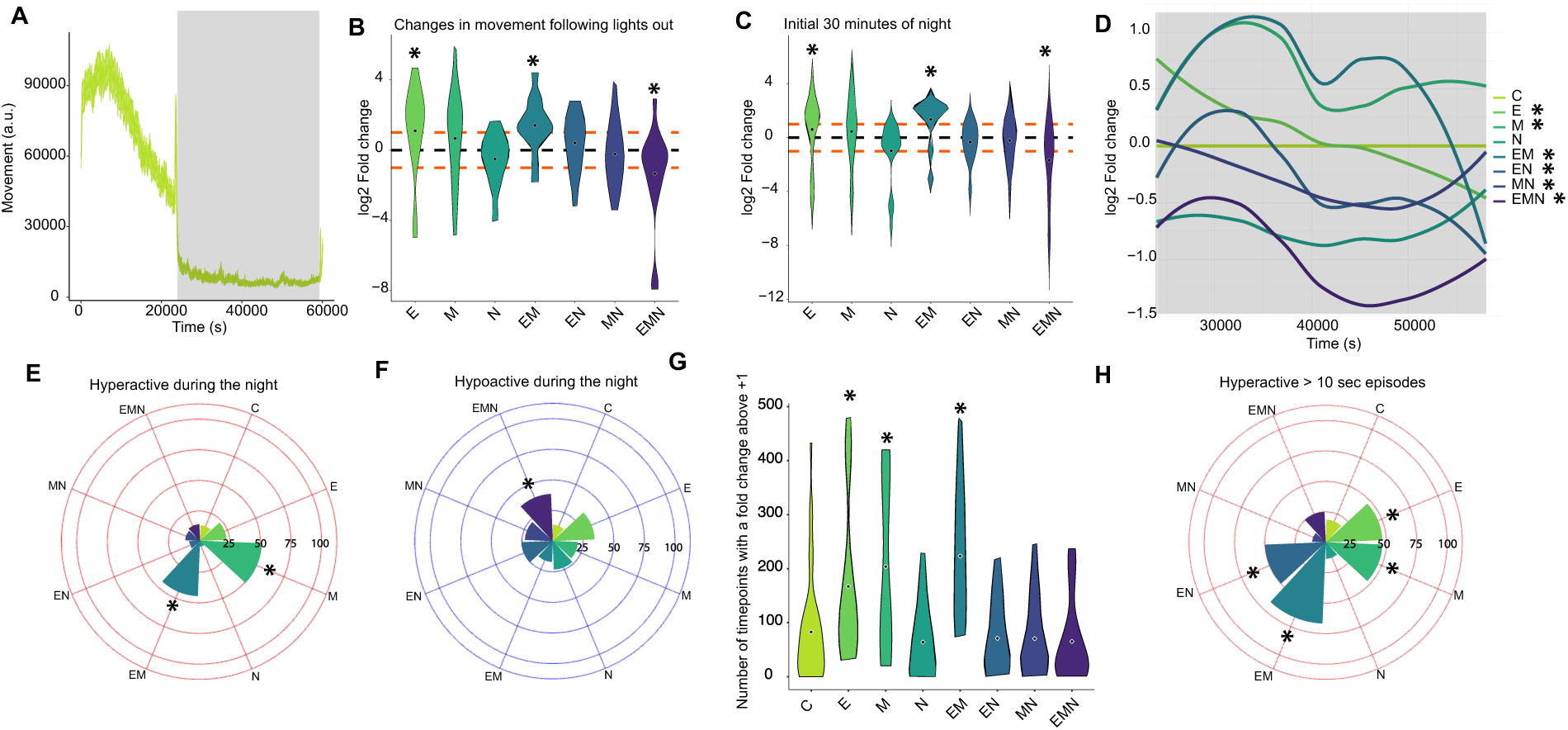
Sleeping pattern in 8 dpf larvae prenatally exposed. **A**. Average movement of control larvae across the sleep test protocol. Movement was calculated using pixel difference. **B.** We calculated the difference between the last 15 minutes of light and the first 15 minutes of dark. E and EM-exposed larvae had a greater behavioural response after the light was turned off, while EMN-exposed animals showed a reduced response compared to controls. * = p < 0.05 compared to control animals. *P-*values were calculated using the Mann– Whitney test followed by Benjamini–Hochberg correction for multiple comparisons. **C.** During the initial 30 minutes of the night, E and EM-exposed larvae showed an increased average log2 fold change in movement. In contrast, EMN-exposed larvae showed a hypoactive phenotype. * = p < 0.05 compared to control animals. *P-*values were calculated using the Mann–Whitney test followed by Benjamini–Hochberg correction for multiple comparisons. **D.** To determine whether drug-exposed larvae exhibited different sleeping patterns, we performed a linear mixed-model analysis to assess changes in log2 fold change during the night. This figure represents the average log2 fold change for each condition over time using a smoothing function. The linear mixed-model analysis revealed distinct shifts in activity patterns during the middle of the night, with EM-exposed larvae showing an abrupt decrease in activity around 45000 seconds. * p < 0.05 compared to control animals. *P-*values were obtained using a linear mixed-effects model with condition over time as a fixed effect and subject (larva ID) as a random effect. **E.** The percentage of larvae with an average log2 fold change of greater than 1 across the dark phase was calculated. * = p < 0.05 compared to control animals. P-values were calculated by using a logical regression. M- and EM-exposed larvae were hyperactive during the night. **F**. The percentage of larvae with an average log2 fold change of lower than -1 across the dark phase was calculated. A greater proportion of larvae in the EMN-exposed group had a lower night-time activity than controls. * = p < 0.05 compared to control animals. *P-*values were calculated by using a logical regression. **G**. The number of time points with a log2 fold change greater than 1 was higher in E, M and EM-exposed groups during the night compared to controls. * = p < 0.05 compared to control animals. *P-*values were calculated using the Mann–Whitney test followed by Benjamini–Hochberg correction for multiple comparisons. **H**. The number of episodes in which the log2 fold change exceeded 1 for 10 consecutive frames was calculated per larva throughout the night. Then, the proportion of animals in each group with more such events than the control average was determined. A higher proportion of E-, M-, EM-, and EN-exposed animals displayed these consecutive hyperactive episodes compared to controls. For all experiments, n ≥ 24 per condition. The violin plot displays the data in log2 fold change relative to control animals. The black dashed line indicates the average of control larvae (0), while the orange lines represent a log2 fold change of +1 and -1. A log2 fold change of +1 represents a twofold increase in movement, while -1 indicates a twofold decrease compared to control animals. This experiment employed a between-subjects design.

We then quantified the light-to-dark transition by comparing activity in the last 15 minutes of light with the first 15 minutes of darkness. E and EM-exposed animals remained more active than controls, whereas EMN larvae showed a rapid drop in movement. N-exposed larvae displayed a similar, though non-significant, trend (**Fig.4B**). Extending the analysis to the first 30 minutes of dark confirmed these patterns: E- and EM-exposed larvae remained hyperactive, while EMN-exposed animals were hypoactive (**Fig.4C**). Although this assay does not directly measure sleep onset, the persistence of sustained activity in the dark in certain drug-exposed groups suggests a delayed transition to rest compared to controls.

By using a linear mixed-effects model, we characterized the temporal dynamics of larval activity across the night. Most groups showed a gradual decline in activity, but the magnitude and direction varied substantially. E-exposed larvae exhibited the steepest reduction, while EM larvae initially displayed marked hyperactivity that progressively shifted toward hypoactivity later in the night. M-exposed larvae maintained sustained hyperactivity, whereas EN and MN groups became progressively less active than controls. EMN larvae were already hypoactive at the start of the night, and this difference grew more pronounced over time **(Fig.4D)**.

Further evidence came from the fact that 48% of M-exposed and 42% of EM-exposed animals showed at least twice the activity level of controls, whereas 38% of EMN larvae showed half the activity level. **(Fig.4E-F).** To investigate sleep fragmentation, we quantified high-activity bursts (log2 fold change greater than 1) and consecutive hyperactive bouts (>10 minutes with a log2 fold change greater than 1), which we interpret as reflecting brief awakenings and nighttime restlessness. Moreover, E-, M-, and EM-exposed larvae exhibited more frequent high-activity bursts and extended hyperactive bouts than controls **(Fig. 4G).** While EN larvae did not show more bursts overall, half displayed more prolonged hyperactive episodes **(Fig. 4H)**. Conversely, EMN larvae exhibited more consecutive hypoactive bouts, with similar though more minor effects in EN and E groups (**Supp.Fig.7)**. Notably, both E- and EN-exposed larvae were overrepresented among individuals showing alternating hyperactive and hypoactive bouts, suggesting fragmented sleep. As with sensorimotor responses, variance analysis revealed greater inter-individual variability during the night across all groups, except EN (**Supp. Fig. 8A**).

During the extended light phase before night, E-exposed larvae were hyperactive, while EMN animals were hypoactive (**Supp.Fig.8B-C**). During the initial 15 minutes of night N, MN, and EMN-exposed larvae were significantly hypoactive (**Supp.Fig.8D**). For the remainder of the light phase, E-exposed animals were hyperactive, as said before. In contrast, N, EN and EMN groups showed reductions in activity (**Supp.Fig.8E)**. Variability before the night was also higher in most drug-exposed groups, except for N and EM (**Supp.Fig.8F)**. When we measured the variance, we noticed that it was dynamic over the duration of the experiment. For instance, variance over time peaked around the middle of the night in all conditions, but occurred earlier in controls, again suggesting altered temporal dynamics in drug-exposed animals (**Supp.Fig.9A**).

The same protocol was repeated at 14 dpf. Control animals displayed activity patterns comparable to those at 8 dpf, with activity increasing during the day and decreasing steadily at night (**Fig.5A**). Across conditions, average traces were broadly similar (**Supp.Fig.10**). However, M-exposed larvae showed an early-night hyperactivity peak, with 50% displaying elevated activity in the first 30 minutes (**Fig.5B**). Across the night, most groups declined in activity, but E-exposed larvae showed the strongest decrease, while EM-exposed animals increased their activity, which is the opposite of their earlier trend **(Fig.5C)**. Distribution analyses revealed that 33 % of E- and EM-exposed animals were at least twice as active as controls **(Fig.5D)**, while 42 % of M- and 33% of EMN-exposed larvae had an average log2 fold change of -1, suggesting a quiescent state **(Fig.5E)**. Hyperactivity metrics showed that the E, M, and EM groups displayed more frequent activity bursts than the controls **(Fig.5F)**. Interestingly, despite this, 53 % of M-exposed larvae also displayed more prolonged hypoactive bouts, again suggesting disrupted sleep (**Fig.5G-H).** While variance peaked mid-night as at 8 dpf (**Supp.Fig.9B)**, patterns differed across treatments: E, M, and EM-exposed larvae showed elevated variance, while N and EN groups showed reduced variance (**Supp.Fig.11A)**. During the extended light phase, 28% of E- and 50% EM-exposed larvae were hyperactive, while 45% of MN animals were hypoactive (**Supp.Fig.11B-C)**. In the first 15 minutes before lights out, E and EM-exposed larvae were significantly more active, while EN and EMN animals were hypoactive (**Supp.Fig.11D)**. For the remainder of the light period, both E and EM larvae maintained higher activity, while N and MN groups exhibited greater variability (**Supp.Fig.11E)**.

**Figure 5:**
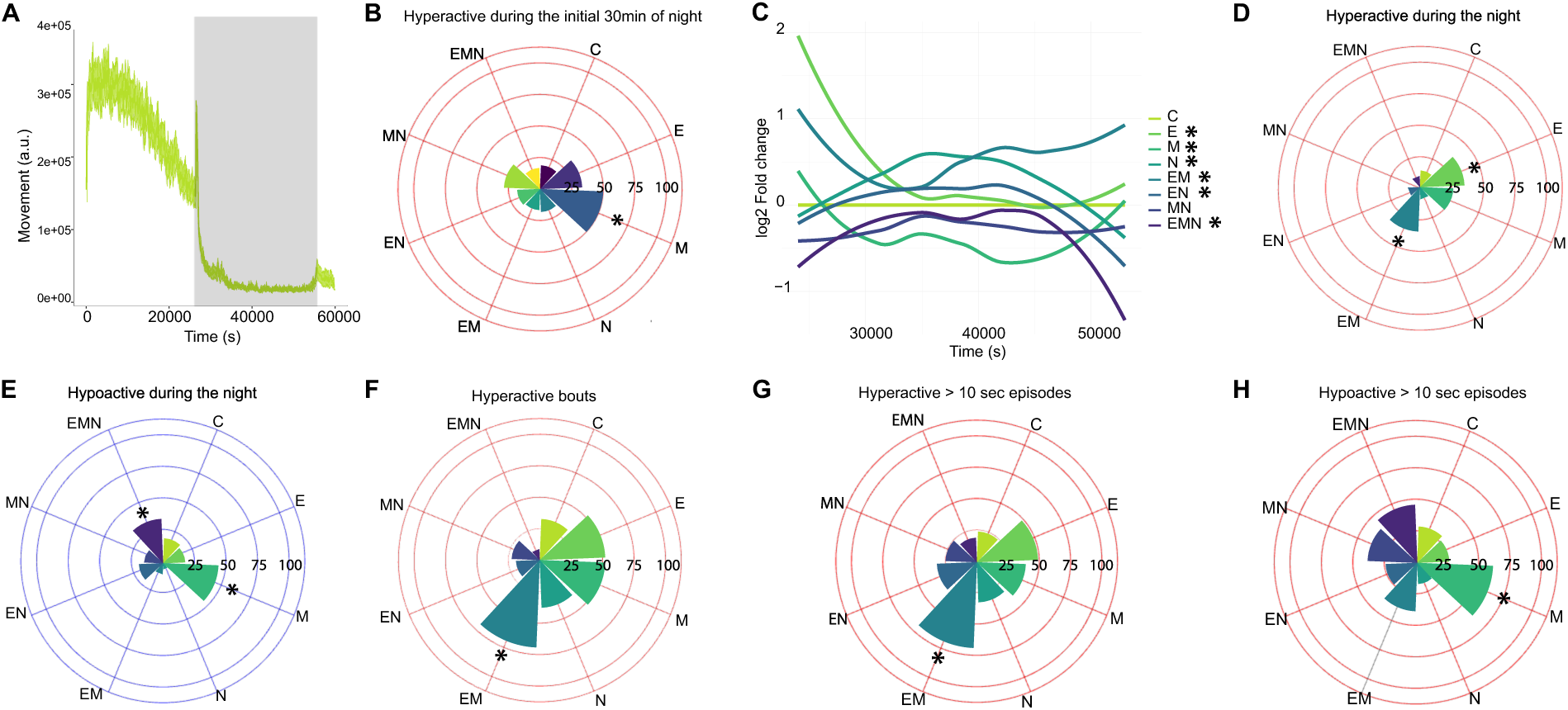
Sleeping pattern in 14 dpf larvae prenatally exposed. **A.** Average movement of control larvae across the sleep test protocol. Movement was calculated using pixel difference. **B.** The percentage of larvae with an average log2 fold change of greater than 1 during the initial 30 minutes of the night was calculated and was higher in the M group. * = p < 0.05 compared to controls. *P-*values were calculated by using a logical regression. **C**. To determine whether drug-exposed larvae exhibited different sleeping patterns during the night, we performed a linear mixed-model analysis to assess changes in log2 fold change. This figure represents the average log2 fold change for each condition over time using a smoothing function. The linear mixed-model analysis revealed that most groups exhibited a decrease in log2 fold change, particularly in E-exposed larvae, whereas EM animals showed the opposite pattern. * p < 0.05 compared to control animals. *P-*values were obtained using a linear mixed-effects model with condition over time as a fixed effect and subject (larva ID) as a random effect. **D.** The percentage of larvae with an average log2 fold change greater than 1 over the entire night was calculated and was higher in the E and EM groups.* = p < 0.05 compared to control animals. *P-*values were calculated using logistic regression. **E.** The percentage of larvae with an average log2 fold change lower than -1 over the entire night was calculated and was higher in the M and EMN groups. * = p < 0.05 compared to control animals. P-values were calculated using logistic regression. **F.** The number of time points at which each larva showed a log2 fold change greater than 1 was calculated. Animals in the EM-exposed group exhibited a higher number of hyperactive events compared to the controls. * = p < 0.05 compared to control animals. *P-*values were calculated using logistic regression. **G.** The number of episodes in which log2 fold change exceeded 1 for 10 consecutive frames was calculated per larva throughout the night. Then, the proportion of animals in each group with more such events than the control average was determined. A higher proportion of EM-exposed animals displayed these consecutive hyperactive episodes compared to controls. **H.** The number of episodes in which log2 fold change was smaller than -1 for 10 consecutive frames was calculated per larva throughout the night. Then, the proportion of animals in each group with more such events than the control average was determined. M-exposed animals had a higher number of larvae showing more consecutive hyperactive events than control animals. For all experiments, n ≥ 24 per condition. The violin plot displays the data in log2 fold change relative to control animals. The black dashed line indicates the average of control larvae (0), while the orange lines represent a log2 fold change of +1 and -1. A log2 fold change of +1 represents a twofold increase in movement, while -1 indicates a twofold decrease compared to control animals. This experiment was conducted using a between-subjects design.

In summary, prenatal drug exposure disrupted sleep-like behaviour at both 8 and 14 dpf. While effects were generally stronger at 8 dpf, treatment-specific patterns persisted. E- and M-exposed animals remained hyperactive, EMN larvae were consistently hypoactive, and several groups showed fragmented nighttime activity. Together, these findings suggest that prenatal exposure alters both the intensity and structure of sleep-wake cycles, with developmental changes over time that may reflect compensatory or maladaptive adaptations of the maturing brain.

### 3.5 Different light-dark preference following prenatal drug exposure

A common outcome of early-life stress is heightened anxiety during both childhood and adolescence(Buckingham-Howes et al. 2012; Juruena et al. 2020). In zebrafish, light-dark preference assays are widely used to assess anxiety-like behaviour as animals can freely choose between a covered (dark) and an uncovered (light) zone. We recently adapted this paradigm for use in zebrafish larvae, which at this developmental stage naturally prefer illuminated environments. Our protocol consisted of 10 minutes of white light, followed by 3 minutes of strobe light as a stressor and a final 10 minutes of white light for recovery. Movement was quantified within the light zone. Rather than measuring time spent in each zone, we focused on locomotor activity in the light area, since reduced activity may indicate either freezing or avoidance (by hiding in the dark zone), both of which reflect stress-related responses.

At 8 dpf, control animals showed a transient reduction in movement during the strobe phase, consistent with freezing or avoidance, and then recovered to baseline before gradually increasing movement (**Fig.6A**). In contrast, over half of the animals exposed to E, M, EM or EMN displayed reduced activity across the entire assay (**Fig.6B**). Linear mixed-effects modelling confirmed a significant decline in log2 fold change over time in several drug-exposed groups, particularly in EN and EMN larvae **(Fig.6C)**.

**Figure 6:**
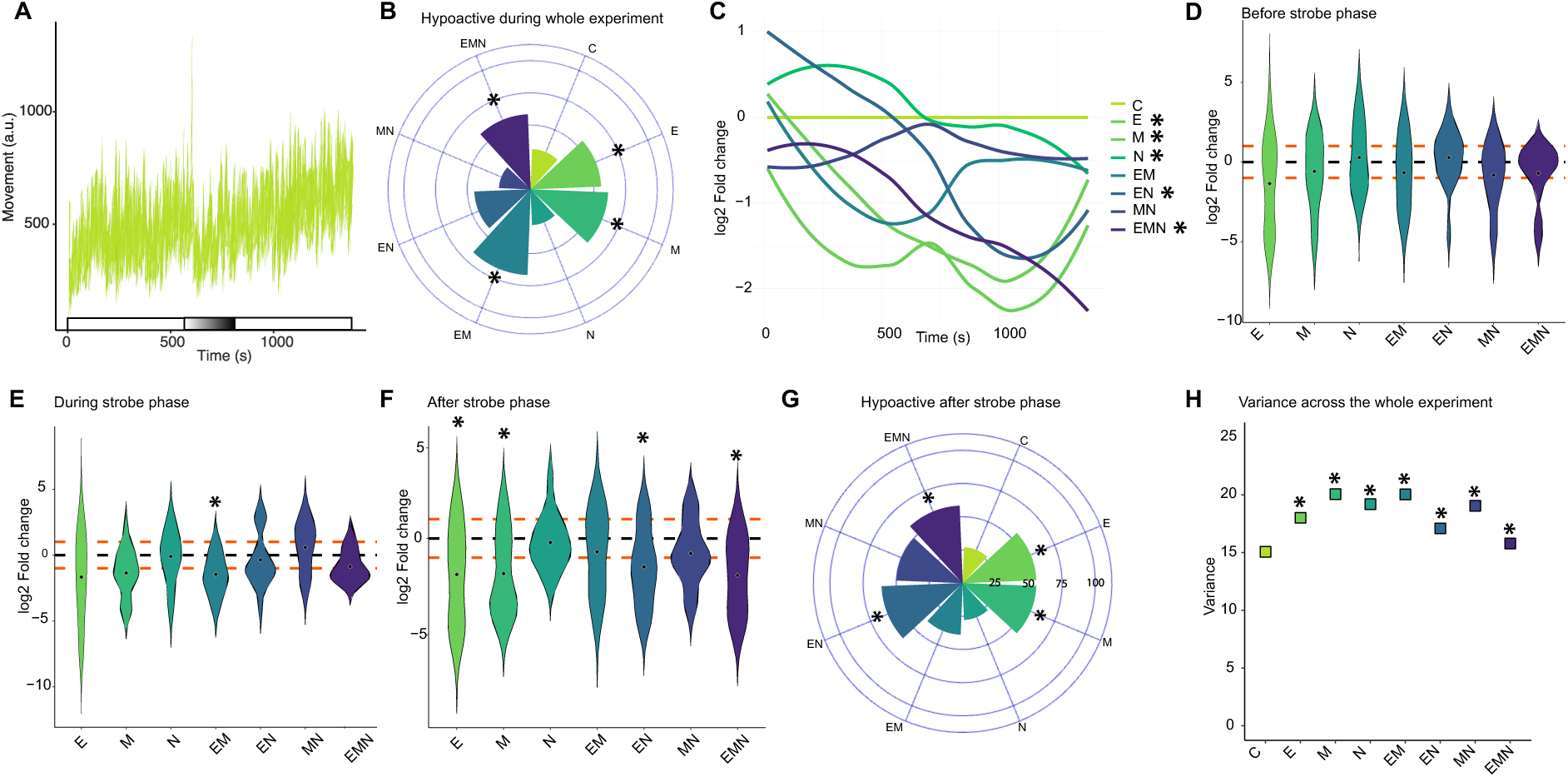
Global activity in response to the dark-light preference test in 8 dpf larvae prenatally exposed. **A**. Average movement of control larvae across the dark-light preference test protocol. Movement was calculated using pixel difference. A reduction in movement can be observed during the strobe light phase. **B.** The percentage of animals with an average log2 fold change lower than -1 across the duration of the entire experiment was calculated and was greater in E, M, EM and EMN groups. * = p < 0.05 compared to control animals. *P-*values were calculated using logistic regression. **C.** To determine the difference between control and drug-exposed larvae throughout the experiment, we performed a linear mixed-model analysis to assess changes in log2 fold change during the strobe phase. This figure represents the average log2 fold change for each condition over time using a smoothing function. The linear mixed-model analysis revealed a reduction in log2 fold change over time in several drug-exposed groups, particularly in the EMN group. * p < 0.05 compared to control animals. *P-*values were obtained using a linear mixed-effects model with condition over time as a fixed effect and subject (larva ID) as a random effect. **D.** During the initial 10 minutes, no significant difference in movement in the light zone in any of the drug-exposed groups was observed. **E.** The strobe phase induced a reduction in movement in the light zone in all conditions, especially in the EM-exposed larvae compared to controls. * = p < 0.05 compared to control animals. *P-*values were calculated using the Mann–Whitney test followed by Benjamini– Hochberg correction for multiple comparisons. **F.** During the last 10 minutes of light following the strobe phase, E, M, EN and EMN-exposed animals showed significantly reduced activity compared to controls. Although not significant, animals in the EM and MN groups also tended to display lower activity levels. * = p < 0.05 compared to control animals. *P-*values were calculated using the Mann–Whitney test followed by Benjamini–Hochberg correction for multiple comparisons. **G.** The percentage of animals with a log2 fold change lower than -1 during the last 10 minutes following the strobe phase was calculated and was greater in E, M, EN, and EMN-exposed groups than controls. * = p < 0.05 compared to control animals. *P-*values were calculated using logistic regression. **H**. Variance was greater in all drug-exposed groups across the entire test duration compared to controls. * = p < 0.05 compared to control animals. Significance was calculated with a Levene test followed by Bonferroni correction for multiple comparisons. For all experiments, n ≥ 24 per condition. The violin plot displays the data in log2 fold change relative to control animals. The black dashed line indicates the average of control larvae (0), while the orange lines represent a log2 fold change of +1 and -1. This experiment was conducted using a between-subjects design.

Differences were minor in the initial light phase and not statistically significant (**Fig.6D**). During the strobe phase, only EM larvae deviated, showing a significant decrease in activity (**Fig.6E**). The strongest effects emerged after the strobe light stimuli: larvae from the E, M, EN, and EMN groups were significantly less active than controls (**Fig.6F**), with 61% of EN and 58% of EMN animals exhibiting reduced locomotion (**Fig.6G**). This persistent suppression of activity suggests a failure to recover from the stressor, as animals continue to avoid the light or remain immobile. As in other assays, drug-exposed groups also displayed greater inter-individual variability, particularly in the single-drug conditions **(Fig.6H).**

At 14 dpf, controls again showed a sharp reduction in movement during the strobe phase (**Fig.7A**). Still, drug-exposed groups displayed more widespread impairments across the entire test, especially under polydrug exposures. Up to 75% of MN and 67% of EMN larvae exhibited less than half the activity of controls **(Fig.7B)**. Log2 fold change analysis revealed a general decline in activity over time in all conditions except MN, which maintained consistent activity, though still below control levels (**Fig.7C)**. No significant differences in movement were detected in the pre-strobe phase (**Fig.7D)**. During the strobe phase, 61% of N- and 67% of MN-exposed animals showed reduced activity (**Fig.7E)**. During the final phase, more than half of larvae from all groups except N exhibited log2 fold change values below -1 (**Fig.7F)**. As with other test, variability was again higher in drug-exposed groups (**Fig.7G**).

**Figure 7:**
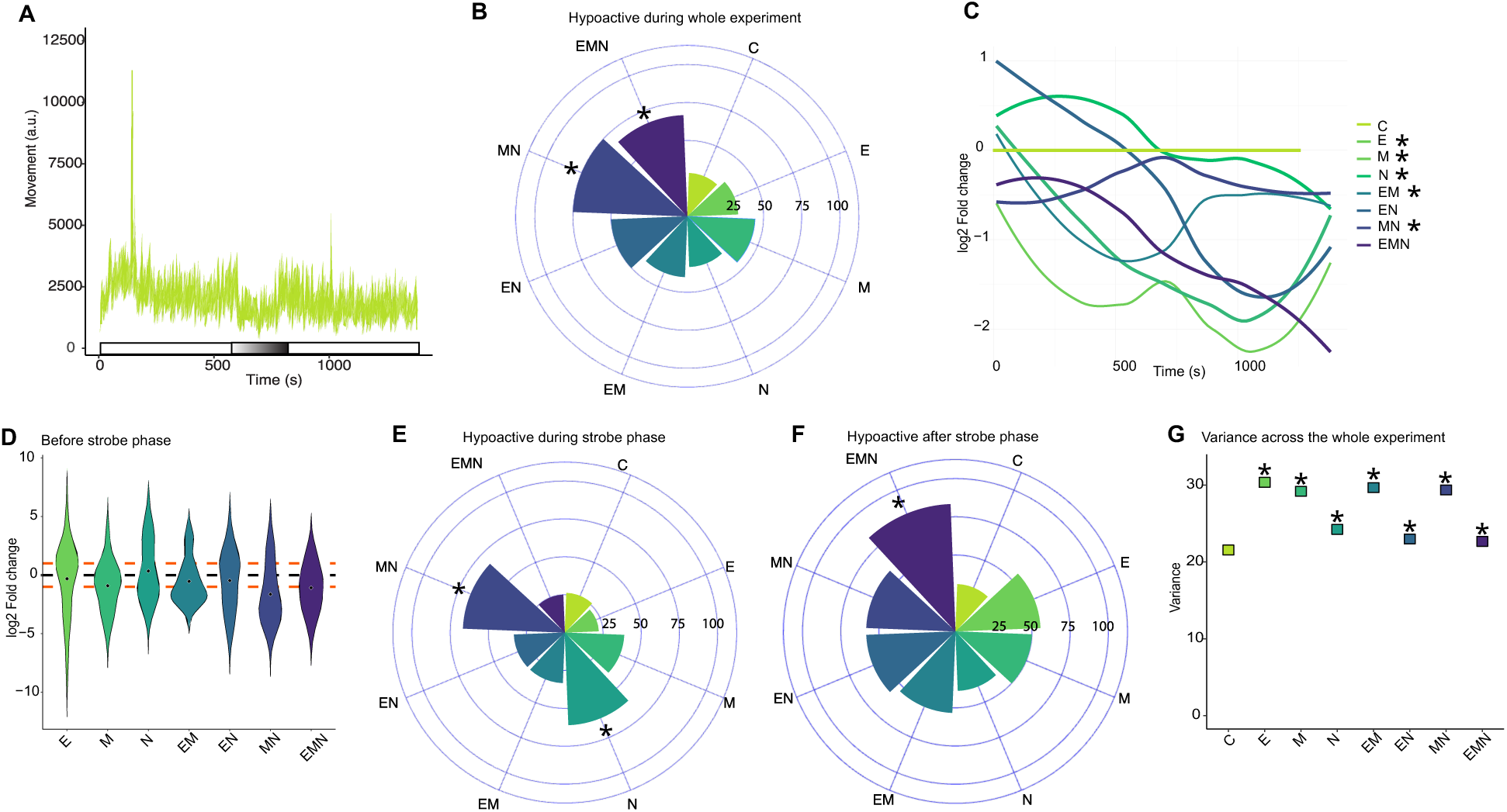
Global activity in response to dark-light preference test in 14 dpf larvae prenatally exposed. **A**. Average movement of control larvae across the dark-light preference test protocol. Movement was calculated using pixel difference. A reduction in movement can be observed during the strobe light phase as at 8 dpf. **B.** The percentage of animals with an average log2 fold change lower than -1 across the duration of the entire experiment is greater in MN and EMN-exposed animals. * = p < 0.05 compared to control animals. *P-*values were calculated using logistic regression. **C.** To determine the difference between control and drug-exposed larvae throughout the experiment, we performed a linear mixed-model analysis to assess changes in log2 fold change variation over time. This figure represents the average log2 fold change for each condition over time using a smoothing function. The linear mixed-model analysis revealed a general reduction in activity over time in all conditions except MN, which maintained consistent activity below control levels.* p < 0.05 compared to control animals. *P-*values were obtained using a linear mixed-effects model with condition over time as a fixed effect and subject (larva ID) as a random effect. **D.** During the initial 10 minutes, there was no significant difference in movement in the light zone in any of the drug-exposed groups. **E.** During the strobe phase, the proportion of animals with a log2 fold change lower than -1 was greater in N and MN-exposed groups. * = p < 0.05 compared to control animals. * = p < 0.05 compared to control animals. *P-*values were calculated using logistic regression. **F.** After the strobe phase, all drug-exposed groups, except for the N-exposed group had at least 50% of larvae with an average log2 fold changer lower than -1 indicating a strong aversion for the light side of the plate. * = p < 0.05 compared to control animals. *P-*values were calculated using logistic regression. **G**. Variance was greater in all drug-exposed groups across the entire test duration compared to controls. * = p < 0.05 compared to control animals. Significance was calculated with a Levene test followed by Bonferroni correction for multiple comparisons. For all experiments, n ≥ 24 per condition. The violin plot displays the data in log2 fold change relative to control animals. The black dashed line indicates the average of control larvae (0), while the orange lines represent a log2 fold change of +1 and -1. A log2 fold change of +1 represents a twofold increase in movement, while -1 indicates a twofold decrease compared to control animals. This experiment was conducted using a between-subjects design.

Together, these results suggest that drug-exposed larvae exhibit altered responses in the light-dark preference assay, particularly when subjected to a stressor. While we do not observe an apparent increase in anxiety-like behaviour, the consistent reduction in activity in the light zone, especially following the strobe light-induced stress, points to altered stress responses. Notably, these effects were more apparent at 8 dpf, while greater variability at 14 dpf among exposed groups may have masked some differences.

### 3.6 Prenatal drug exposure alters thermosensation

Following our initial behavioural assays, we extended our characterization to a different sensory modality by implementing a thermosensation assay. Adapted from the rodent hot plate paradigm, this test measures nociceptive-like responses to a rapid increase in temperature, which in zebrafish larvae manifests as increased locomotion. In controls, this response was robust (**Fig.8A)**, and individual traces revealed drug-specific alterations, most notably enhanced activity in M-exposed groups (**Supp.Fig.12)**

**Figure 8:**
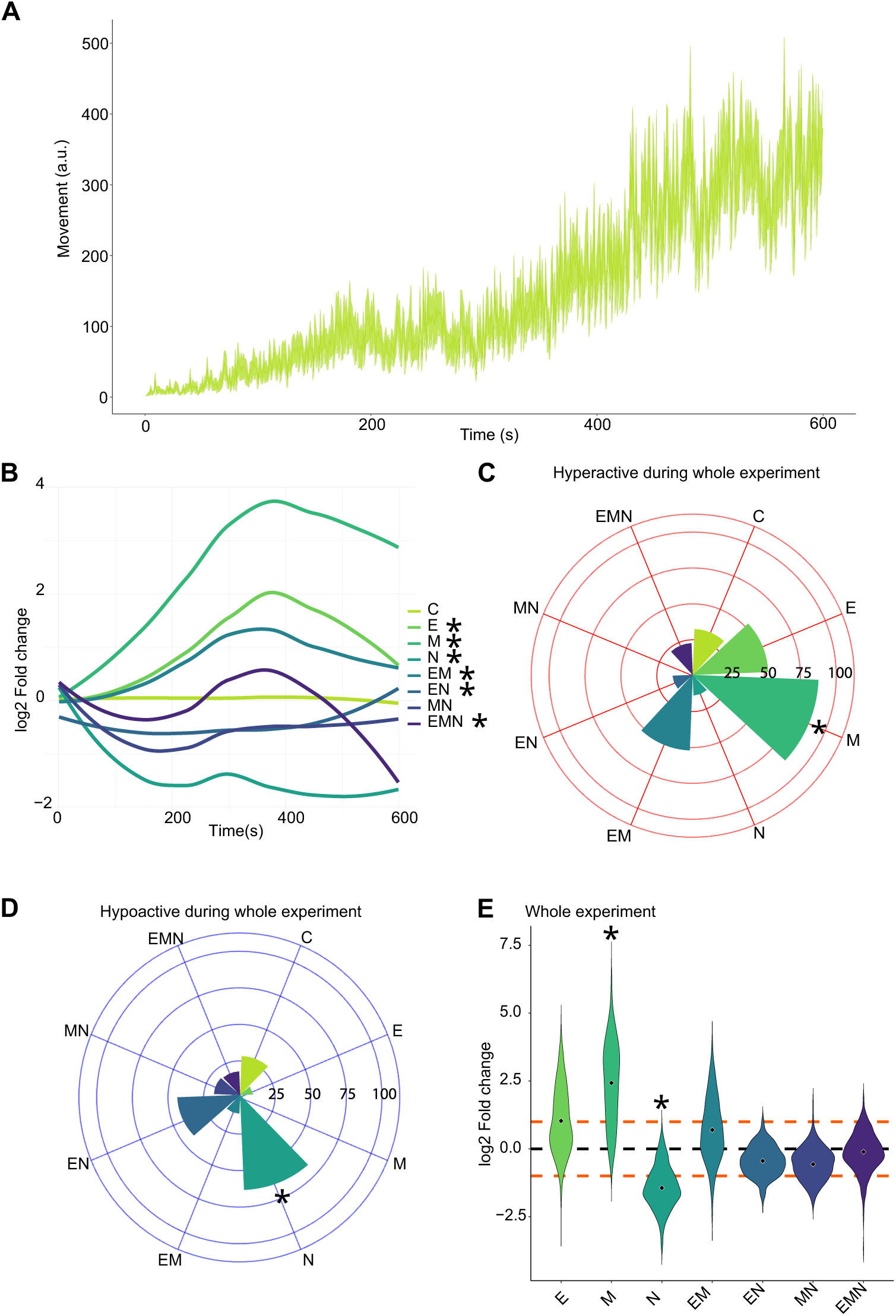
Thermal response to heat in 8 dpf larvae prenatally exposed. **A.** Average locomotor responses over time of control larvae throughout the thermosensation assay. **B.** To determine the difference between control and drug-exposed larvae throughout the experiment, we performed a linear mixed-model analysis to assess changes in log2 fold change variation over time. This figure represents the average log2 fold change for each condition over time using a smoothing function. The linear mixed-model analysis revealed larvae in the E, M and EM groups showed an enhanced average fold-change over time, while N-exposed groups showed a clear reduction in responsiveness. * p < 0.05 compared to control animals. *P-*values were obtained using a linear mixed-effects model with condition over time as a fixed effect and subject (larva ID) as a random effect. **C.** The percentage of larvae with an average log2 fold change of greater than 1 across the whole duration of the experiment was exceeding 75% in the M-exposed group. * = p < 0.05 compared to control animals. * = p < 0.05 compared to control animals. *P-*values were calculated using logistic regression. **D.** The percentage of larvae with an average log2 fold change of lower than -1 across the whole-duration of the experiment was calculated and was the highest in the N-exposed group. **E.** The average log2 fold change over the duration of the experiment was significantly higher in M-exposed larvae compared to controls, while the opposite is observed in the N-exposed animals. * = p < 0.05 compared to controls. *P-*values were calculated using the Mann–Whitney test followed by Benjamini–Hochberg correction for multiple comparisons. For all experiments, n ≥ 24 per condition. The violin plot displays the data in log2 fold change relative to control animals. The black dashed line indicates the average of control larvae (0), while the orange lines represent a log2 fold change of +1 and -1. A log2 fold change of +1 represents a twofold increase in movement, while -1 indicates a twofold decrease compared to control animals. This experiment was conducted using a between-subjects design.

At 8 dpf, drug-exposed animals displayed distinct response profiles compared to controls. Notably, larvae in the E, M and EM groups showed enhanced log2 fold changes, evident as early as 60 seconds after the test’s onset. In contrast, N-exposed groups showed an apparent reduction in responsiveness **(Fig.8B)**. Morphine produced the most pronounced effect, with 82% of individuals exhibiting more than a 2-fold increase in activity compared to controls (**Fig.8C**). Conversely, 64 % of N-exposed larvae showed a strong decrease in response (**Fig.8D**). Comparing the end of the assay to its onset further highlighted these trends: M-exposed animals reached a 2.5 log2 fold increase, with E- and EM-exposed animals also showing sustained hyperactivity. In contrast, N-, EN, and MN-exposed animals displayed reduced activity relative to controls.

At 14 dpf, controls maintained thermosensory responses comparable to those observed at 8 dpf (**Fig. 9A)**. The drug-exposed groups, however, diverged from the earlier results. E and M-exposed larvae now showed an average 2-fold decrease in activity. In contrast, animals from the triple-exposed group displayed a 2-fold increase suggesting heightened sensitivity to heat in the latter (**Fig.9B and Supp. Fig.13**). Distribution analysis revealed that multi-drug-exposed groups contained a higher proportion of hyper-responsive individuals: 100% of EMN, 54% of EN and 50% of EM larvae had log2 fold change values above 1 (**Fig.9C)**. In contrast, reduced responsiveness dominated in single-drug groups, with 64% of M- and 50% of E-exposed animals showing log2 fold change values below –1, indicating a reduced reactivity to water temperature changes (**Fig.9D**). These group-level shifts were also reflected in mean activity: M-exposed animals showed a significant decrease, while EMN-exposed animals exhibited a significant increase (**Fig.9E).**

**Figure 9:**
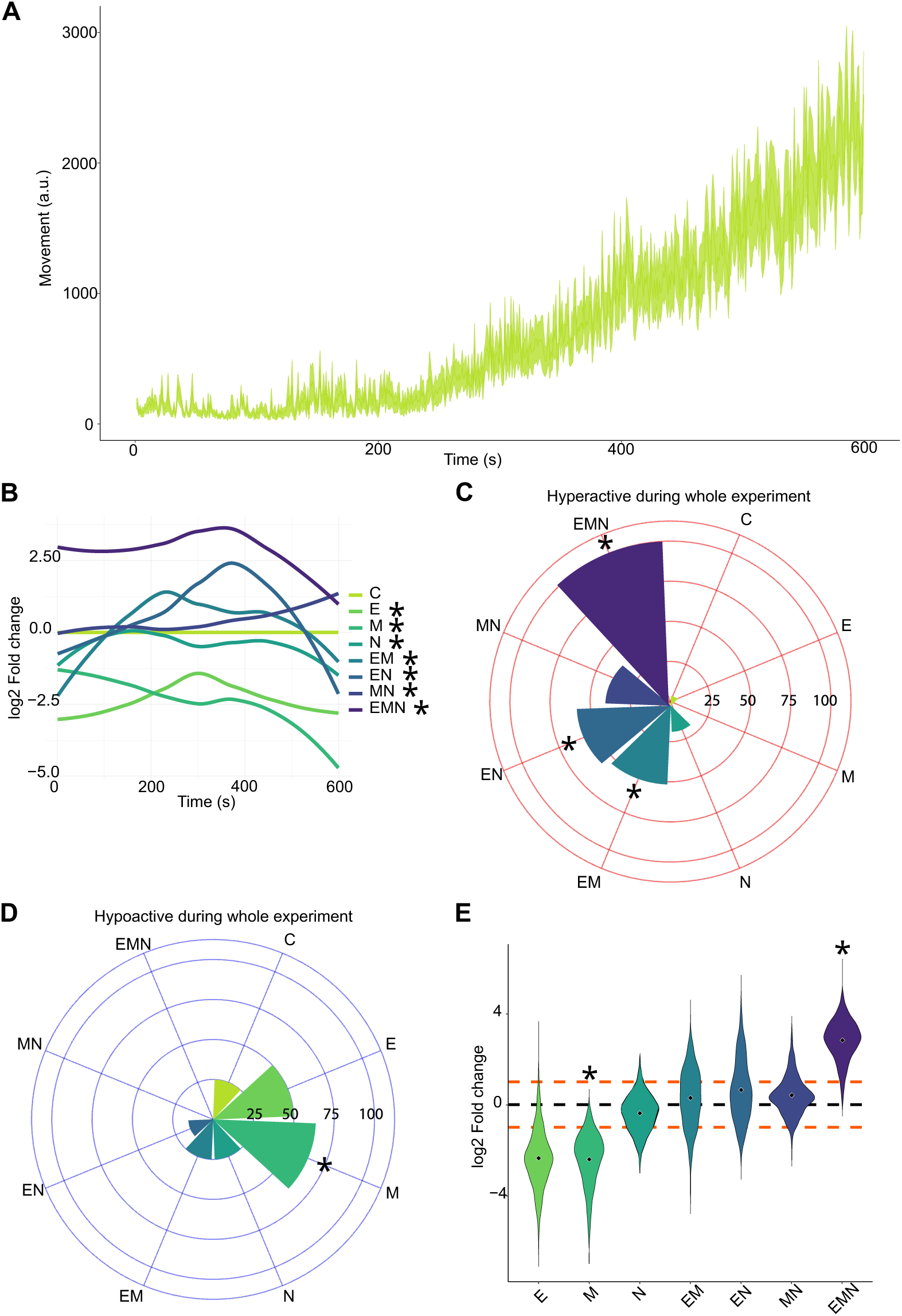
Thermal response to heat in 14 dpf larvae prenatally exposed. **A.** Average locomotor responses over time of control larvae throughout the thermosensation assay. **B.** To determine the difference between control and drug-exposed larvae throughout the experiment, we performed a linear mixed-model analysis to assess changes in log2 fold change variation over time. This figure represents the average log2 fold change for each condition over time using a smoothing function. The linear mixed-model analysis revealed that larvae exposed to ethanol or morphine showed decreased activity compared to controls, whereas triple-exposed animals showed the opposite trend over time, suggesting heightened sensitivity to heat in the latter. * p < 0.05 compared to control animals. *P-*values were obtained using a linear mixed-effects model with condition over time as a fixed effect and subject (larva ID) as a random effect. **C.** Nearly all EMN- and more than half of EM- and EN-exposed larvae presented an average log2 fold change greater than 1 over the total duration of the thermosensation assay. * = p < 0.05 compared to control animals. *P-*values were calculated using logistic regression. **D.** The percentage of larvae with an average log2 fold change of lower than -1 across the whole-duration of the experiment was calculated and was the highest in the M-exposed group. **E.** The average fold change over the duration of the experiment was significantly higher in EMN-exposed larvae compared to controls, while the opposite trend was observed in the M-exposed animals. * = p < 0.05 compared to control animals. *P-*values were calculated using the Mann–Whitney test followed by Benjamini–Hochberg correction for multiple comparisons. For all experiments, n ≥ 24 per condition. The violin plot displays the data in log2 fold change relative to control animals. The black dashed line indicates the average of control larvae (0), while the orange lines represent a log2 fold change of +1 and -1. A log2 fold change of +1 represents a twofold increase in movement, while -1 indicates a twofold decrease compared to control animals. This experiment employed a between-subjects design.

In summary, morphine exposure promoted strong heat-evoked activity at 8 dpf but suppressed responses by 14 dpf, whereas EMN exposure produced the opposite effect. These findings highlight how single versus combined drug exposures can produce divergent, even opposing, outcomes that evolve during development.

### 3.7 Alterations in social interaction dynamics

Zebrafish begin to exhibit social behaviours during the second and third weeks of development, with sociability becoming more robust by 21 dpf(Dreosti et al. 2015). To assess the impact of drug exposure on social preference, we used an established two-chamber paradigm where larvae could choose between proximity to conspecifics or an empty compartment (**Supp.Fig.14A)**. A social index was calculated for each larva and expressed as log2 fold change relative to its clutch-matched control. Although group averages did not differ significantly from controls, distinct trends emerged: Animals exposed to E, N or the MN combination showed elevated mean indices, suggesting increased sociability, whereas M- and EM-exposed larvae showed reduced mean indices (**Supp.Fig.14B)**. Specifically, 81 % of E- and 67 % of N-exposed larvae exhibited a social index more than twice that of controls (**Supp.Fig.14C)**. Conversely, 50% of M-, 40% EM- and 75% of EN-exposed larvae showed a social index at least two-fold lower (**Supp.Fig.14D)**. Distributional analysis further underscored these shifts. For example, despite no significant change in mean, over 75% of E-exposed animals clustered in the “highly social” category, an effect that diminished when ethanol was combined. Notably, no “non-social” individuals were detected in the MN group, suggesting a potential balancing or protective effect of this combination **(Fig.10E)**.

Overall, while social preference was not significantly altered at the group level, drug-exposed larvae exhibited marked shifts in individual sociability. These findings highlight the complex and sometimes opposing effects of prenatal drug exposure on early social development.

## 5. Discussion

Polydrug exposure during pregnancy remains a significant public health issue, impacting millions of children worldwide. While decades of research have explored the effects of individual drugs such as alcohol, nicotine, and opioids, growing clinical evidence suggests that single-drug exposure is more the exception than the rule(Jarlenski et al. 2017). Despite this, the biological consequences of prenatal polydrug exposure are still not well understood. In this study, we used the zebrafish model to conduct a behavioural analysis of the short- and long-term effects of developmental exposure to ethanol, nicotine, and morphine, three commonly used substances that affect different neurochemical pathways.

Our goal was to determine whether drug combinations would produce specific effects, rather than merely additive or opposing effects. Our findings demonstrate that prenatal exposure to drugs of abuse induces complex and dynamic behavioural changes in zebrafish larvae.

### Behavioral outcomes

Overall, our findings highlight that polydrug exposure can result in synergistic, antagonistic, or compensatory interactions, depending on the receptor targets and timing of action. For example, while exposure to a single drug does not lead to an increase in mortality, combining nicotine and ethanol enhances toxicity, suggesting either synergistic or additive effects. On the other hand, at the behavioural level, the combination of ethanol and nicotine leads to moderate differences that could be consistent with an antagonistic interaction or the trigger of compensatory mechanisms.

Across multiple assays, some phenotypes were consistent. In the light-dark preference test, most drug-exposed larvae showed reduced movement in the light zone, especially following stress (strobe light). This pattern suggests heightened anxiety-like behaviour, a common outcome of prenatal drug exposure. Other phenotypes, such as altered sleep and nighttime activity, also parallel clinical reports of disrupted sleep in infants prenatally exposed to alcohol or opioids(Kenner and D’Apolito 1997; Chandler-Mather et al. 2021).

Drug-specific behavioural patterns were also evident. Ethanol and morphine tended to induce hyperactivity, while nicotine consistently produced hypoactivity, which could reflect their respective effects on GABAergic, glutamatergic, cholinergic, dopaminergic, and opioid systems(Dani and Bertrand 2007; Trescot et al. 2008, 2017). Using hierarchical clustering of responses during the MSA revealed that some drug treatments share behavioural patterns, such as the combination of ethanol and morphine, which tends to produce additive-like effects in 8 dpf larvae. Our findings also support the inclusion of a broad spectrum of behavioural assays to present a more comprehensive understanding. For example, at 14 dpf, temperature-based assays demonstrated more pronounced differences than light-based assays, thus emphasizing the importance of multimodal testing.

### Developmental dynamic

By performing the same assay on the same group of animals at different development stages, we were able to observe how behavioural outcomes change over time. One notable feature of our data is the shift in phenotype severity and variability as development progresses. At 8 days post-fertilization (dpf), drug-induced behavioural differences were robust and relatively consistent across animals. By 14 dpf, these effects became more variable and, in some cases, diminished or reversed. For example, morphine-induced hyperactivity at 8 dpf was replaced by hypoactivity at 14 dpf, while the triple-exposed group became hyperactive only at a later stage. This suggests that early behavioural impacts are not static and may evolve due to compensatory plasticity, delayed toxicity, or ongoing brain development. The reversal of the effect of prenatal morphine exposure on the response to temperature changes is a striking example of these differences. This reversal highlights the dynamic nature of behavioural outcomes throughout development and may indicate adaptive and maladaptive reorganization of sensory-motor circuits.

### Population variability

One key finding we observed is the increased variability in the drug-exposed population. This mirrors findings in humans, where prenatal exposure to drugs leads to a wide range of outcomes, from severe impairments to apparently normal development(Behnke et al. 2013). Clustering analysis reinforced this heterogeneity, identifying subgroups of hyper-and hypoactive individuals within treatments. For example, ethanol, morphine, and their combination often produced hyperactive subtypes, while nicotine-containing conditions were more associated with hypoactivity. Notably, morphine + nicotine did not produce hyperactivity at any stage, suggesting possible antagonism. These findings emphasize the importance of expanding our analysis beyond group-level averages to capture individual differences that may reflect resilience, vulnerability or delayed effects.

### Mechanistic interpretation

The increased toxicity observed in polydrug conditions remains to be determined. Possible mechanisms include neurotoxicity or other biological mechanisms, such as elevated oxidative stress, mitochondrial dysfunction, or neuroinflammation, which have been noted to increase following prenatal drug exposure(Das and Vasudevan 2007; Wang et al. 2022). It is also possible that the combination of ethanol and nicotine leads to altered activation of nicotinic receptors, as they can have a synergistic effect(Engle et al. 2015).

Some of the behavioural differences we observed can provide insights into dysregulated neurological pathways. Some of the effects we observed, such as shifts from hyperactivity to hypoactivity, may reflect developmental changes in the balance of excitatory and inhibitory signals or plasticity. Exposure to substances like ethanol or morphine often results in a hyperactive baseline, possibly due to a balance tilted towards excitation(Basnet et al. 2019). Conversely, nicotine-exposed larvae tend to be hypoactive, indicating increased inhibition. Other behaviours, such as nighttime hyperactivity, are also observed, which could be consistent with an excitatory/inhibitory imbalance or dysregulation of other systems regulating sleep, such as orexin or norepinephrine(Sohal and Rubenstein 2019).

Our findings underscore the importance of further studies to investigate the molecular effects of such exposure and identify causative alterations.

### Limitations

The combination of ethanol and nicotine, along with the EMN group, was associated with higher mortality; therefore, the more moderate behavioural phenotypes observed in those groups could result from survivor bias, as only the most resilient individuals might have survived long enough to be tested. Furthermore, we primarily measured sensorimotor response using a specific behavioural assay, so employing different types of assays could have yielded different results. It is also possible that changes at the molecular level may not be fully captured in our behavioural assays.

## 6. Conclusion

Together, our data show that the effects of prenatal drug exposure are not only drug-specific but also influenced by drug combinations. While some changes emerge early and persist throughout development, other phenotypes continue to evolve over time. These findings are directly relevant to understanding how polydrug use during pregnancy can influence neurodevelopmental trajectories in offspring. Since polydrug use is common in human populations and outcomes can vary significantly even among individuals with similar exposures, future research should aim to identify biomarkers of vulnerability and resilience. This would enable targeted interventions in at-risk populations and help reduce the long-term effects of prenatal drug exposure.

## Data statement

Data and custom tracking algorithms are available upon request.

## Authorship

L.H. Conceptualization, Methodology, Investigation, Formal analysis, Visualization, Writing – Original draft, Writing – Reviewing and editing.

G.D.B Conceptualization, Formal Analysis, Resources, Methodology, Software, Validation, Visualization, Data Curation, Supervision, Funding acquisition, Project Supervision, Writing – Original draft, Writing – Reviewing and editing.

## Acknowledgments

We would like to acknowledge the LARSEM and the animal care facility at CERVO for providing Zebrafish husbandry, laboratory space, and equipment to facilitate portions of this research.

## Abbreviations

dpf: days postfertilization
E: ethanol
fps: frames per second
hpf: hours postfertilization
M: morphine
MSA: multi-stimuli assay
N: nicotine
WT-TL: Wild-type Tübingen long fin

