## Supplemental figures for "Prenatal Polysubstance Exposure Alters Behaviour in Zebrafish Larvae"

### Supplementary data

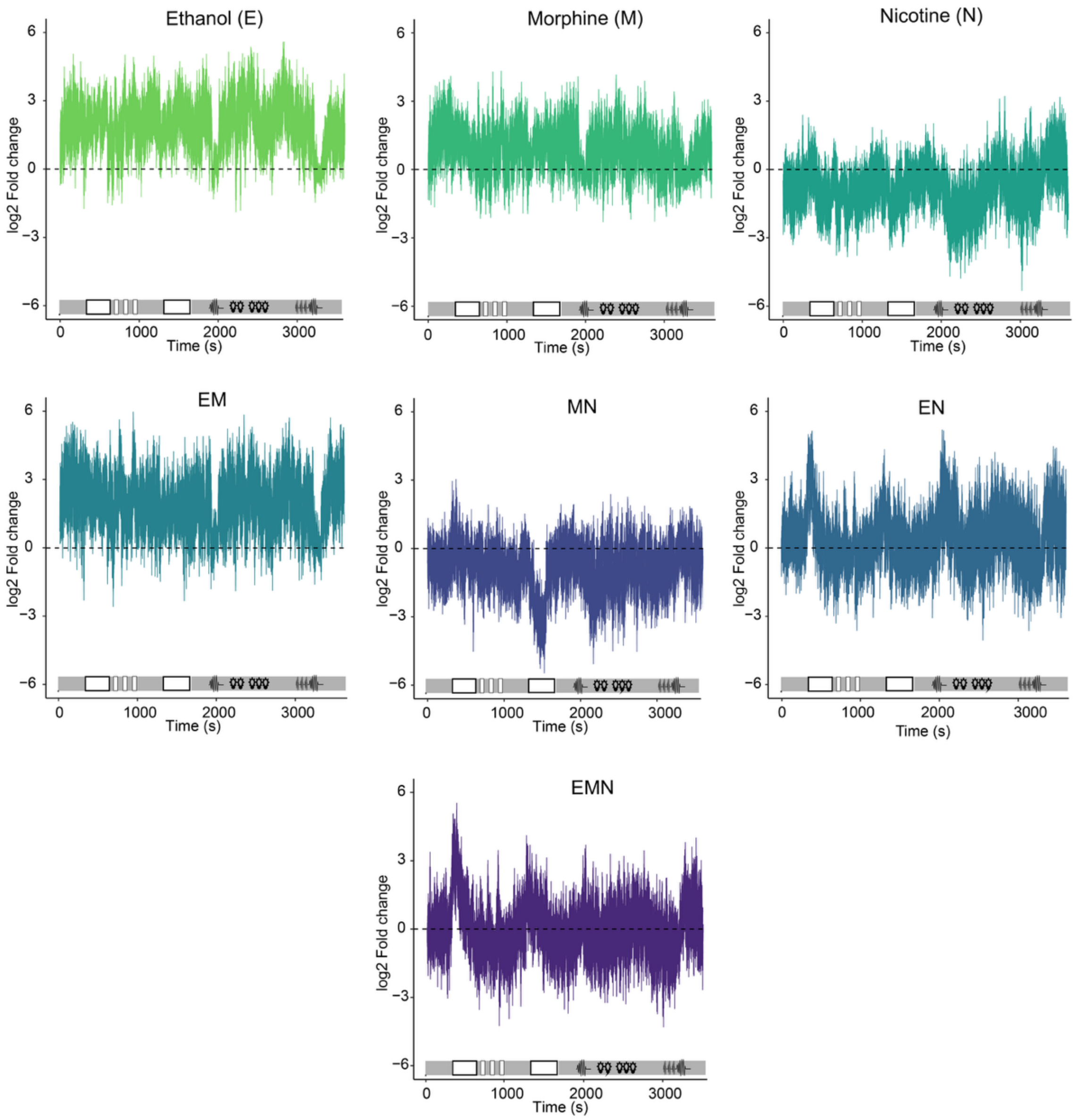


**Supp.Fig.1: Traces of log2 fold change in movement of 8 dpf-larvae across the MSA for each condition relative to controls.**

Average locomotor response of larvae from each group throughout the duration of the MSA. Larvae exposed to E, M, or their combination showed higher activity log2 fold changes compared to controls, while those exposed to N and the MN combination tended to be hypoactive. Animals in the EN and EMN groups exhibited responses similar to those of the controls. $n \geq 24$ per condition

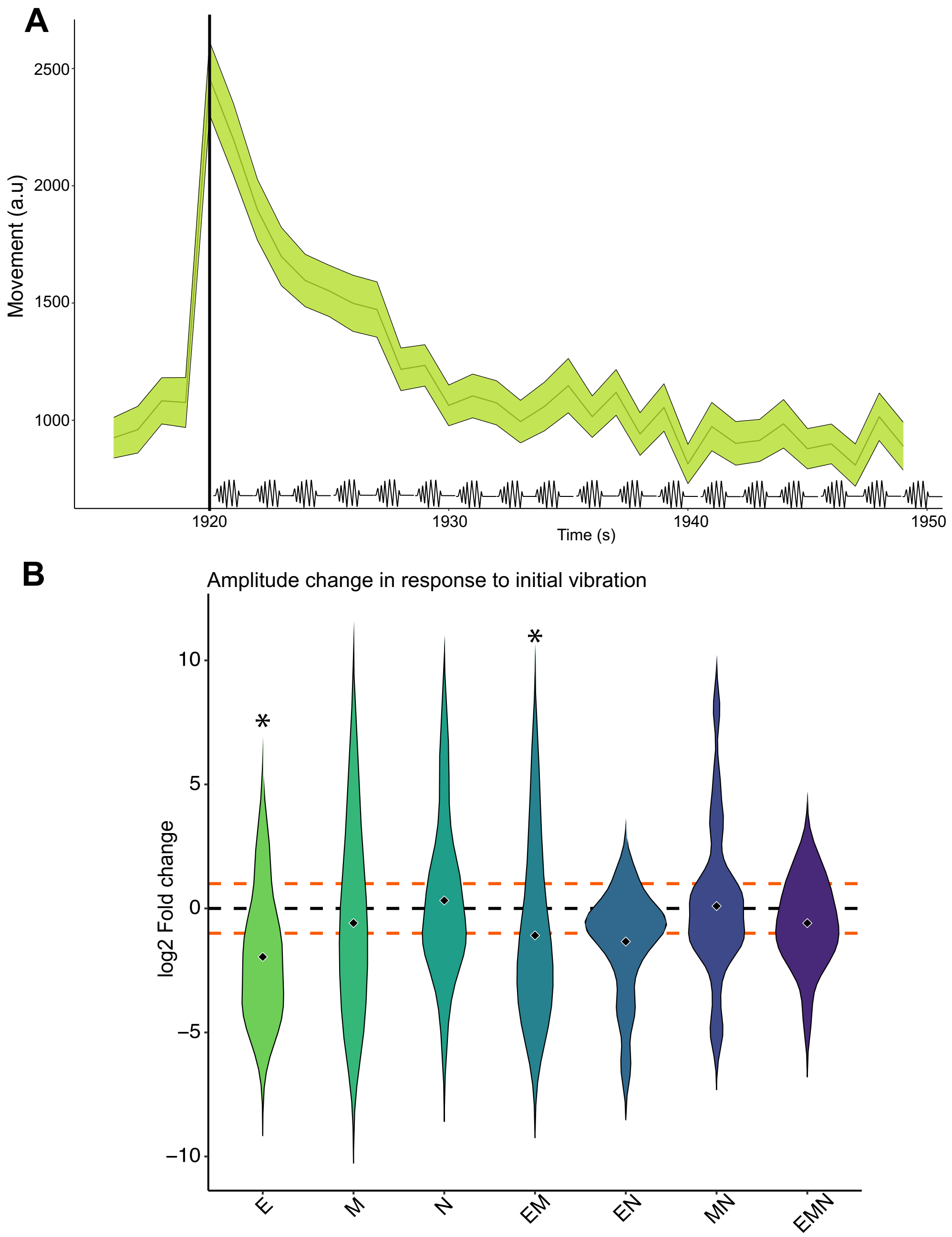


**Supp.Fig.2: Response to repeating vibration during the MSA at 8 dpf.**

**A.** Repeated vibration exposure will lead to habituation and a reduction in response. The black line indicates the initiation of vibration. The green line represents the average response for control animals, and the ribbon shows the average ± SEM. **B.** We calculated the changes in amplitude following the initiation of the vibration sequence. The amplitude was calculated by averaging the first 3 seconds following the first vibration, minus the average of the last 5 seconds before initiation. Animals exposed to ethanol and ethanol-morphine have a lower amplitude than the control. * = $p < 0.05$ compared to control animals. P-values were calculated using the Mann–Whitney test followed by Benjamini–Hochberg correction for multiple comparisons. $n \geq 24$ per condition. This experiment employed a between-subjects design.

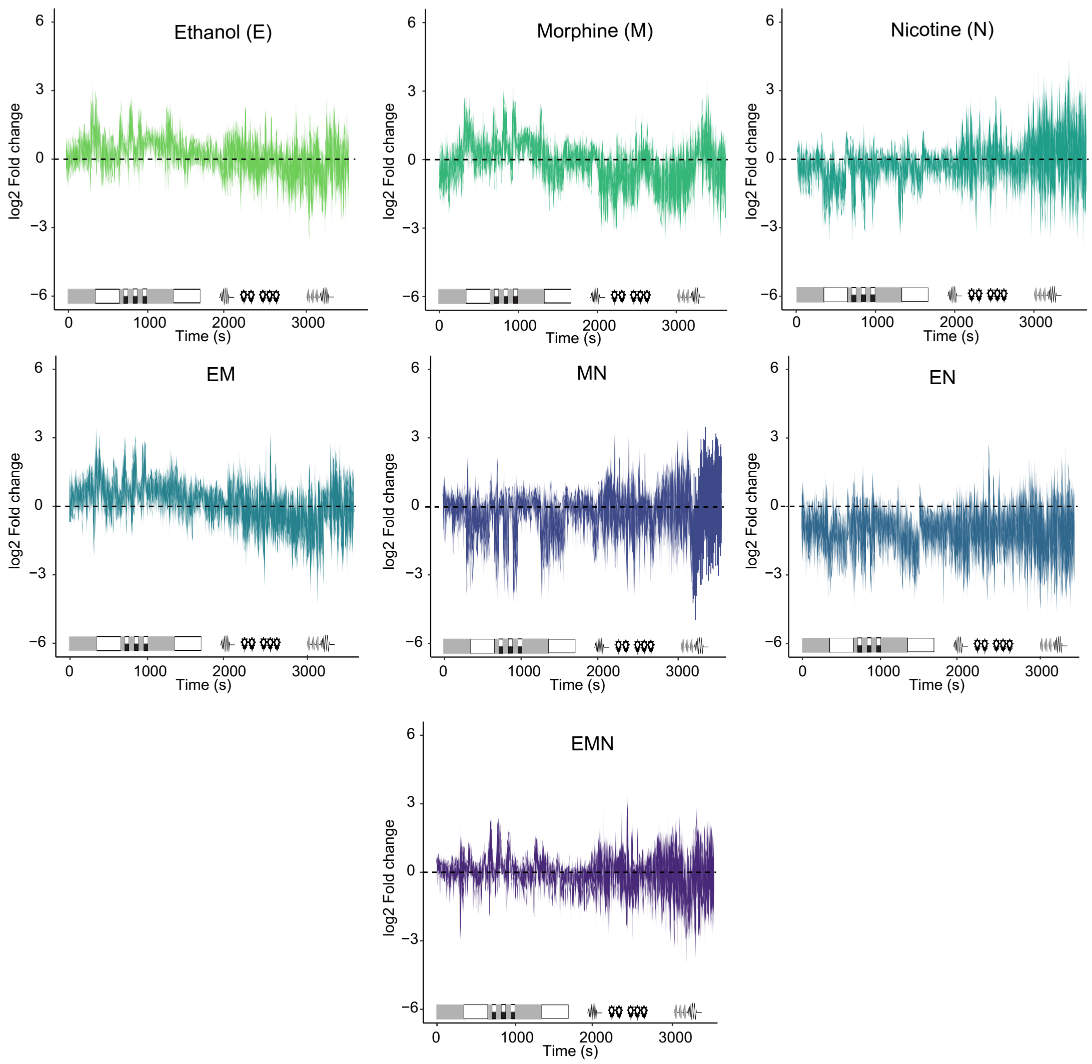


**Supp.Fig.3: Traces of log2 fold change in movement of 14 dpf-larvae across the MSA for each condition relative to controls.**

Average locomotor response of larvae from each group throughout the duration of the MSA. No consistent hyperactive behaviour was observed. Larvae in the EN group tended to be slightly hypoactive, with average log2 fold change below that of controls. $n \geq 24$ per condition.

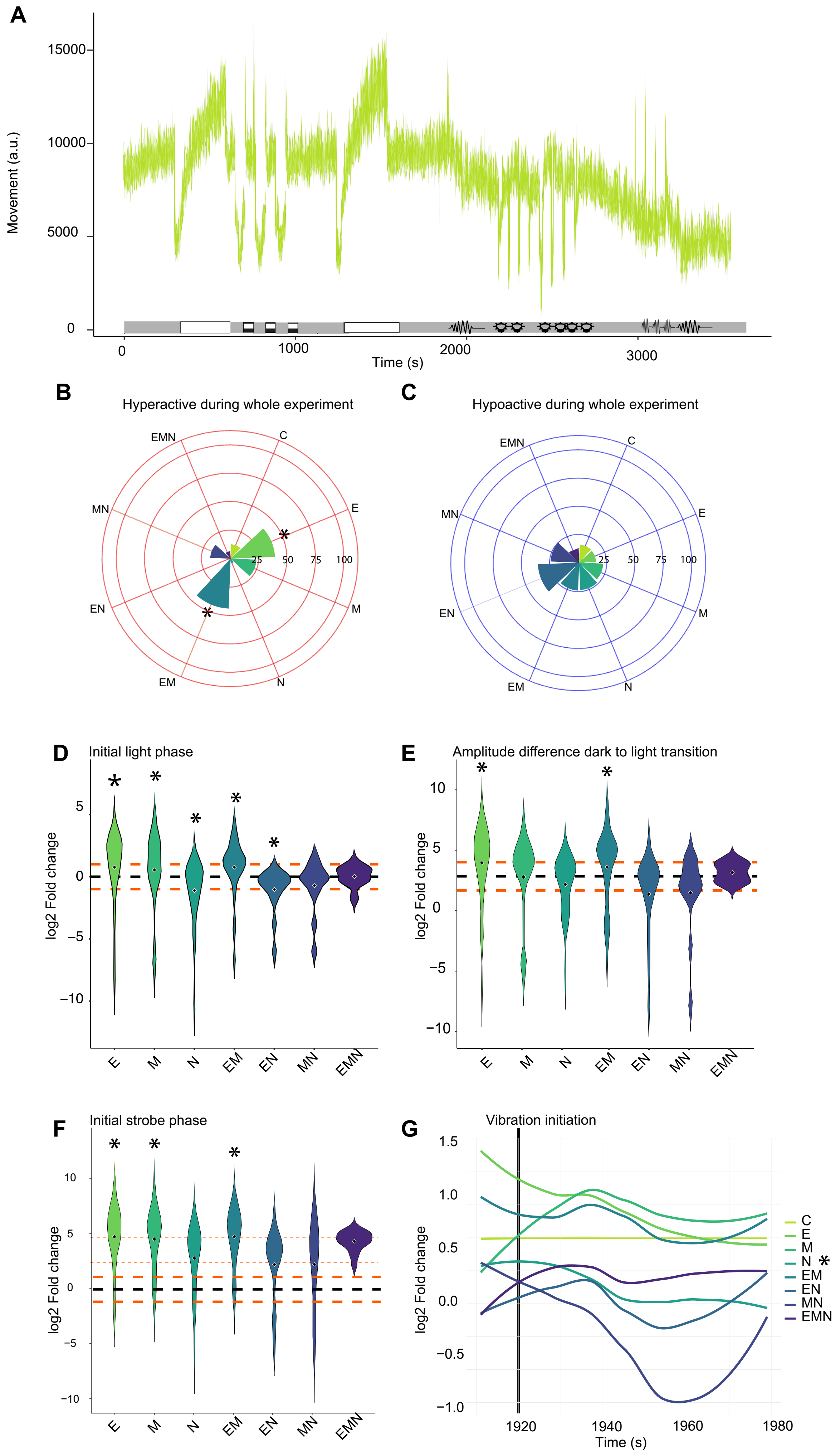

A
Movement (a.u.)
15000
10000
5000
0
0
1000
2000
3000
Time (s)
B
Hyperactive during whole experiment
EMN
C
E
MN
EN
M
EM
N
25
50
75
100
C
Hypoactive during whole experiment
D
Initial light phase
log2 Fold change
E
M
N
EM
EN
MN
EMN
E
Amplitude difference dark to light transition
F
Initial strobe phase
G
Vibration initiation
1.5
1.0
0.5
0.0
−0.5
−1.0
1920
1940
1960
1980
Time (s)
C
E
M
N *
EM
EN
MN
EMN

**Supp.Fig.4: Sensorimotor responses to Multi-Stimuli Assay (MSA) in 14 dpf larvae prenatally exposed.**

**A.** Average locomotor responses of control larvae throughout the duration of the MSA. **B.** The percentage of larvae with an average log2 fold change of greater than 1 across the whole duration of the experiment was calculated. * = $p < 0.05$ compared to control animals. P-values were calculated by using a logistic regression. Nearly half of the EM-exposed animals were hyperactive. **C.** The percentage of larvae with an average log2 fold change of lower than -1 across the whole duration of the experiment was calculated. * = $p < 0.05$ compared to control animals. P-values were calculated by using a logistic regression. Half of the MN group showed hypoactivity. **D.** During the initial light phase, larvae exposed to E, M, or their combination displayed hyperactive behaviour. In contrast, larvae exposed to N, EN or MN were hypoactive. * = $p < 0.05$ compared to control animals. P-values were calculated using the Mann–Whitney test followed by Benjamini–Hochberg correction for multiple comparisons. **E.** Transition from dark to light triggers a high amplitude response in the E and EM groups. We measured the average log2 fold change for the last 5 seconds before the stimulus and the first 3 seconds immediately following the transition. Similar patterns were observed during transitions from dark to light. * = $p < 0.05$ compared to control animals. P-values were calculated using the Mann–Whitney test followed by Benjamini–Hochberg correction for multiple comparisons. **F.** During the total of the three strobe phases, E-, M- and EM- exposed larvae were hyperactive, while N-, EN- and MN- exposed larvae were hypoactive. * = $p < 0.05$ compared to control animals. P-values were calculated using the Mann–Whitney test followed by Benjamini–Hochberg correction for multiple comparisons. **G.** Repeated vibration exposure will lead to habituation and a reduction in response. A linear mixed-model analysis showed that all groups, except those exposed to M, display normal habituation patterns. * $p < 0.05$ compared to control animals. P-values were obtained using a linear mixed-effects model with condition over time as a fixed effect and subject (larva ID) as a random effect. For all experiments, $n \geq 24$ per condition. The violin plot displays the data in log2 fold change relative to control animals. The black dashed line indicates the average of control larvae (0), while the orange lines represent a log2 fold change of +1 and -1. A log2 fold change of +1 represents a twofold increase in movement, while -1 indicates a twofold decrease compared to control animals. This experiment employed a between-subjects design.

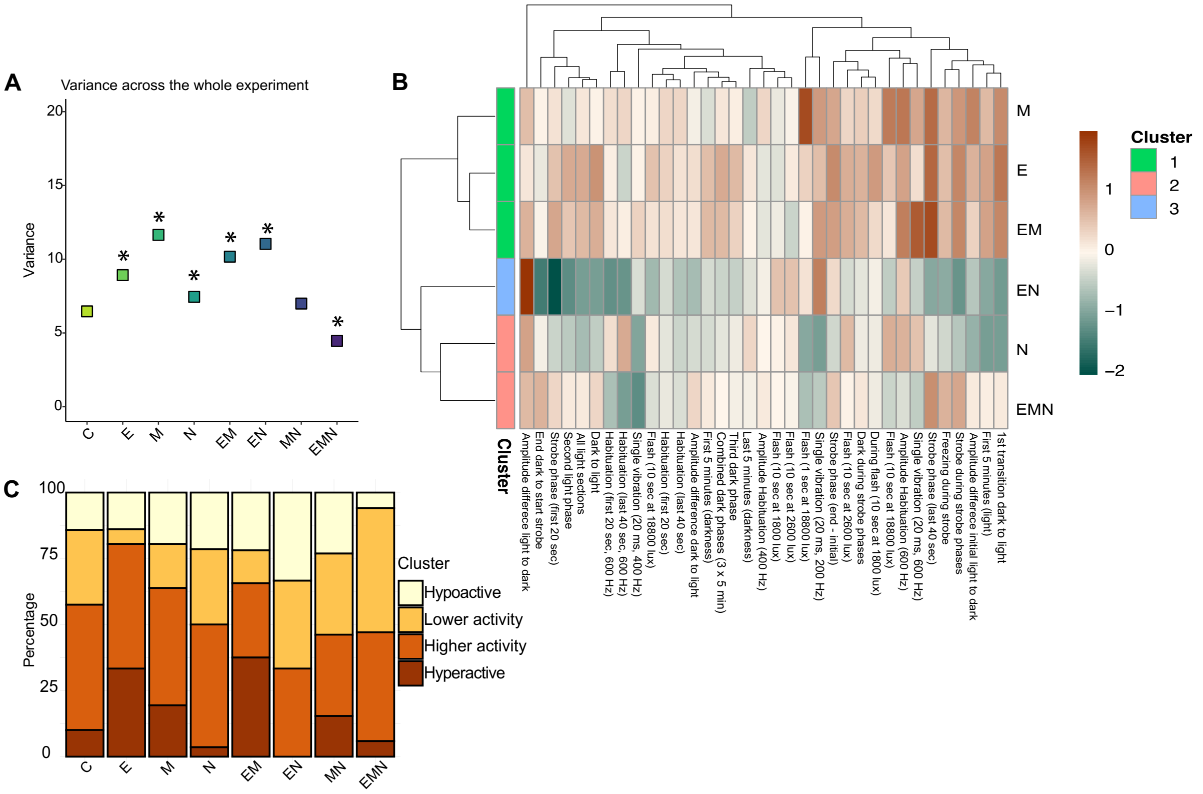


**Supp.Fig.5: Sensorimotor responses to Multi-Stimuli Assay (MSA) in 14 dpf wild-type zebrafish larvae prenatally exposed.**

**A.** Animals from single drug-exposed groups and from the EM and EN groups had a greater variance across the duration of the MSA. * = $p < 0.05$ compared to the control group. Significance was calculated with a Levene test followed by a Bonferroni correction for multiple comparisons. **B.** Manual clustering based on the average log2 fold change for each larva across the duration of the MSA. The categories were separated as follows: Hypoactive log2 fold change lower than -1, Lower activity log2 fold change between -1 and 0, Higher activity log2 fold change between 0 and 1 and Hyperactive log2 fold change greater than 1. Control animals showed a homogenous distribution. No hyperactive individuals were observed in the EN group. **C.** Hierarchical clustering of the different treatments (y-axis) based on the log2 fold change across the various parameters we measured. Heatmap colours indicate the log2 fold change compared to control, where orange and blue reflect increased and decreased responses, respectively. Clusters were then generated based on similar responses. The animals in the morphine-nicotine group showed responses outside the average range for other conditions. This condition was removed from the analysis to facilitate the comparison of the remaining groups. $n \geq 24$ per condition.

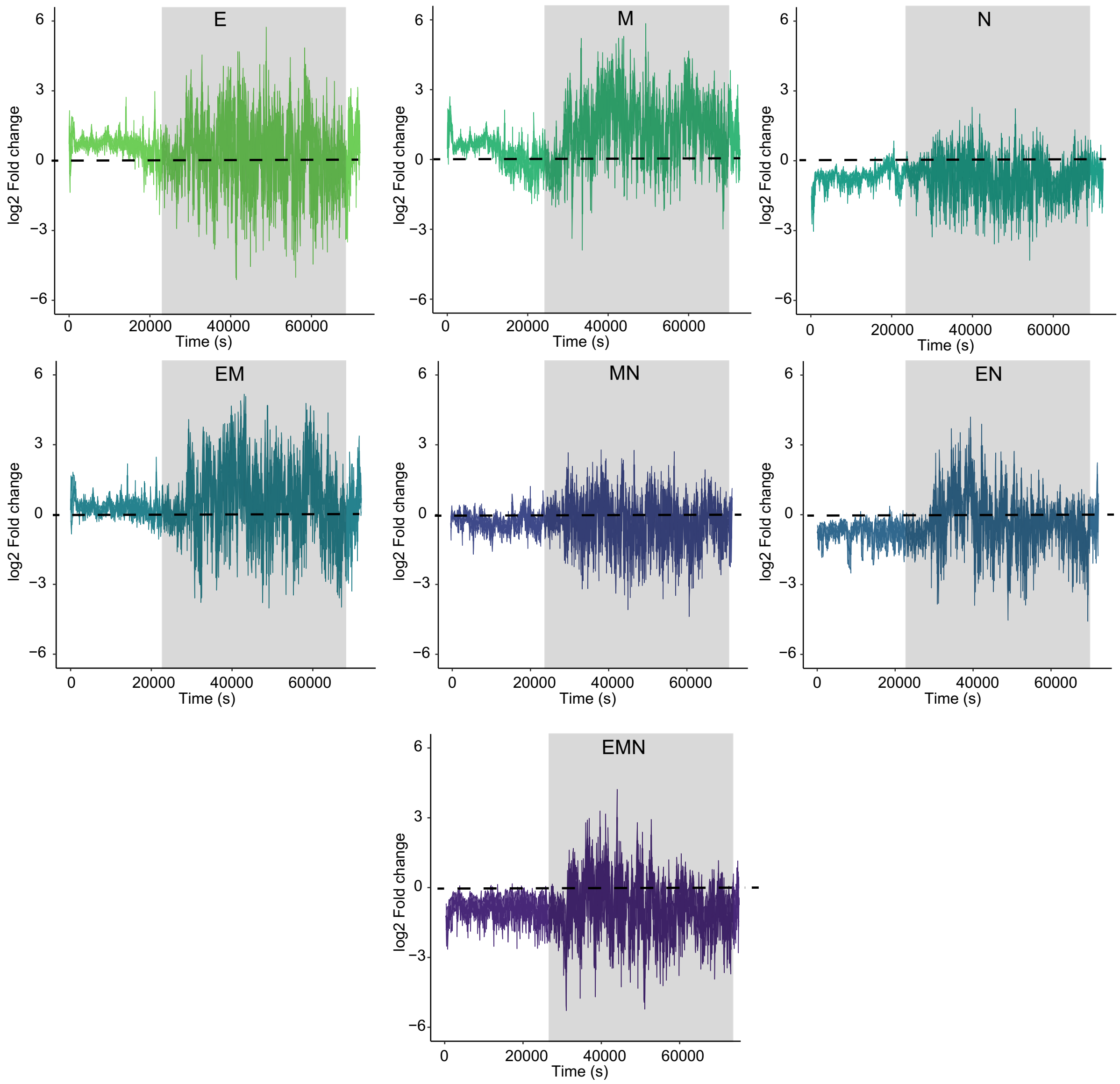


**Supp.Fig.6: Traces of log2 fold change in movement of 8 dpf-larvae throughout the sleep test for each condition relative to controls.**

Average locomotor response of larvae from each group throughout the duration of the sleep test. Larvae exposed to E, M and EM showed increased activity log2 fold changes during the night, along with disrupted sleep-like patterns characterized by irregular fluctuations in activity levels. Larvae from the N, MN, EN and EMN groups also showed altered nighttime activity patterns; however, their average activity log2 fold change remains lower than that of controls, indicating a trend toward hypoactivity. $n \geq 24$ per condition.

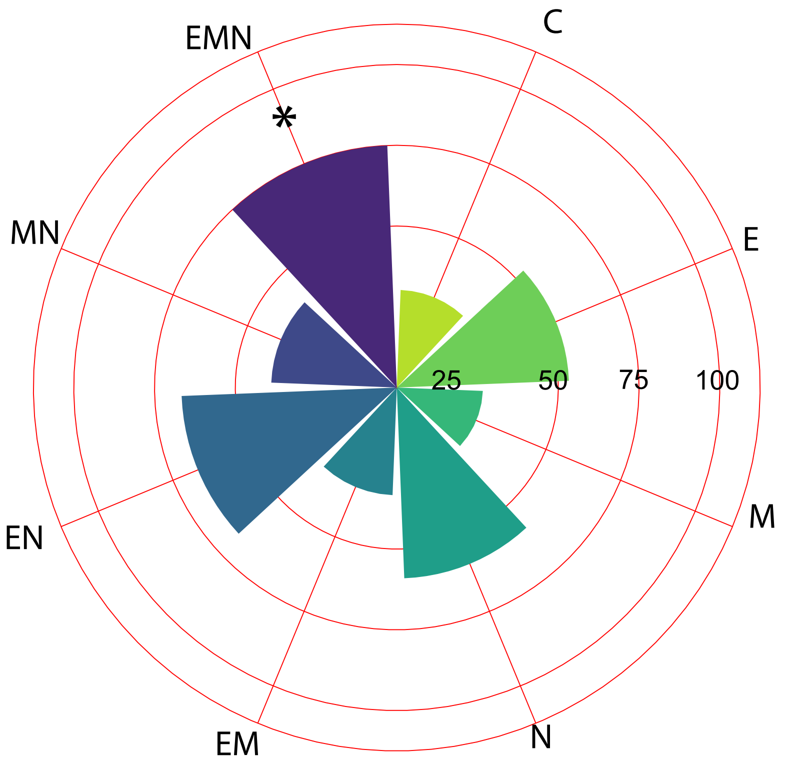


**Supp.Fig.7: Percentage of larvae with an average fold change lower than -1 for 10 consecutive frames across the night.**

The number of times each larva experienced a log2 fold change lower than -1 for 10 consecutive frames was calculated throughout the night. Then, the proportion of animals in each group with a greater number of those events than the average of the control was determined. 75% of EMN-exposed larvae were twice less active than controls. P-values were calculated by using a logistic regression. $n \geq 24$ per condition.

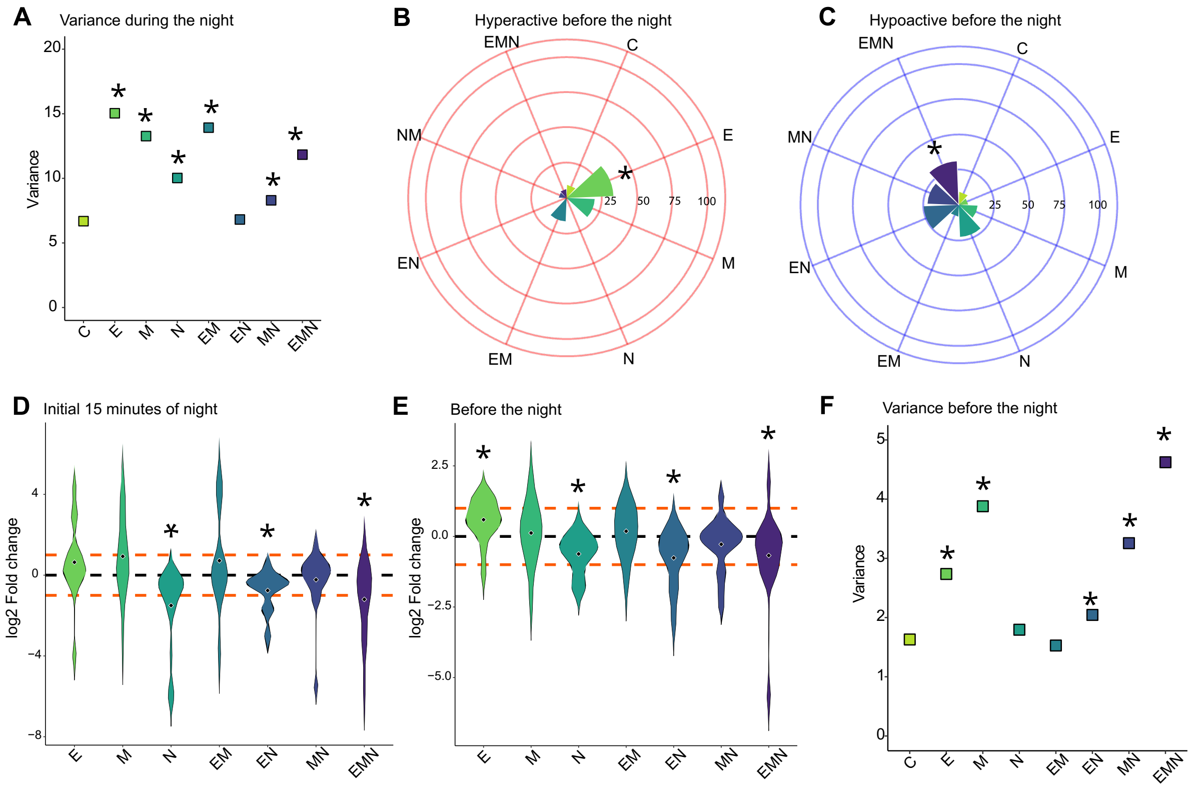


**Supp.Fig.8: Behavioural variability and log2 fold change in movement before and during the night at 8 dpf.**

**A.** Animals from all exposed groups except EN had a greater variance during the night. * = p < 0.05 compared to the control group. Significance was determined using a Levene test followed by a Bonferroni correction for multiple comparisons. **B.** The percentage of larvae with an average log2 fold change of greater than 1 before the dark phase was calculated. * = p < 0.05 compared to control animals. P-values were calculated by using a logistic regression. Over 25% of E-exposed larvae displayed hyperactivity. **C.** The percentage of larvae with an average log2 fold change of lower than -1 before the dark phase was calculated. * = p < 0.05 compared to control animals. P-values were calculated by using a logistic regression. EMN-exposed animals tended to be hypoactive. **D.** During the initial 15 minutes of the night**,** MN and EMN-exposed larvae showed heightened activity, while N-exposed animals exhibited reduced activity. **E.** Before the dark phase, E-exposed animals were more active, while N-, EN- and EMN-exposed groups showed reduced activity. **F.** Animals from the E, M, EN, MN and especially EMN had a greater variance before the night.* = p < 0.05 compared to the control group. Significance was determined using a Levene test followed by a Bonferroni correction for multiple comparisons. For all experiments, n ≥ 24 per condition. The violin plot displays the data in log2 fold change relative to control animals. The black dashed line indicates the average of control larvae (0), while the orange lines represent a log2 fold change of +1 and -1. A log2 fold change of +1 represents a twofold increase in movement, while -1 indicates a twofold decrease compared to control animals. This experiment employed a between-subjects design.

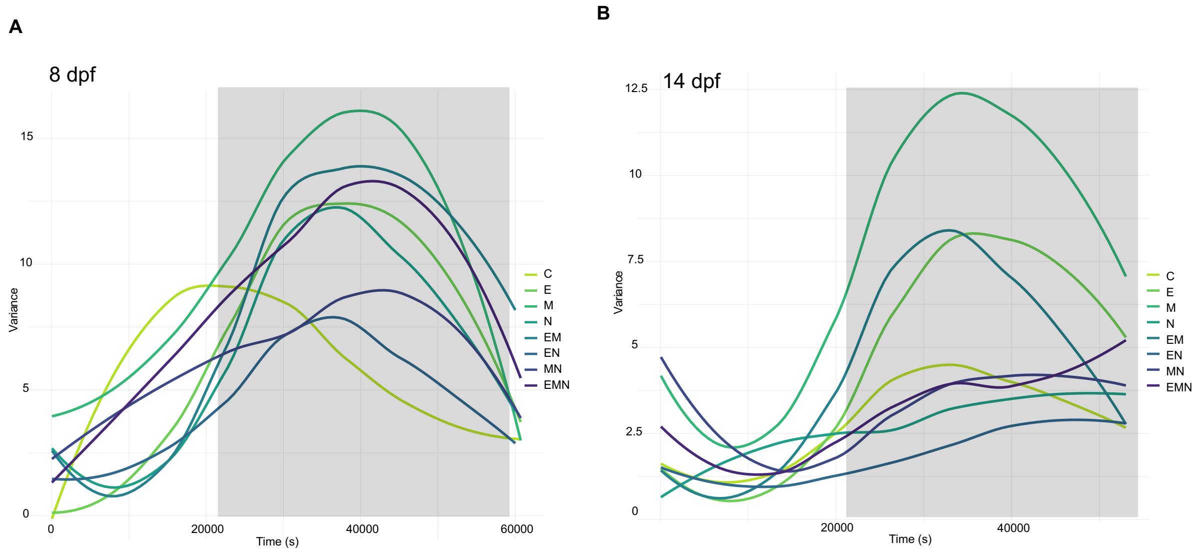


**Supp.Fig.9: Inter-individual variability in movement across the whole sleep test.**

**A**: Variance traces across the entire test at 8 dpf. Before the night phase, drug-exposed animals generally exhibited lower inter-individual variability than controls. However, this trend reversed during the night, with control animals showing the lowest variability. n ≥ 24 per condition. **B:** Variance traces across the entire test at 14 dpf. M-exposed larvae were hyperactive before the onset of the night phase, and this heightened activity persisted throughout the night. In contrast, EN-exposed animals were hypoactive before the night and remained so during the night phase. Traces represent a smoothing of the average variance across all larvae. P-values were obtained using a linear mixed-effects model with condition over time as a fixed effect and subject (larva ID) as a random effect. n ≥ 24 per condition.

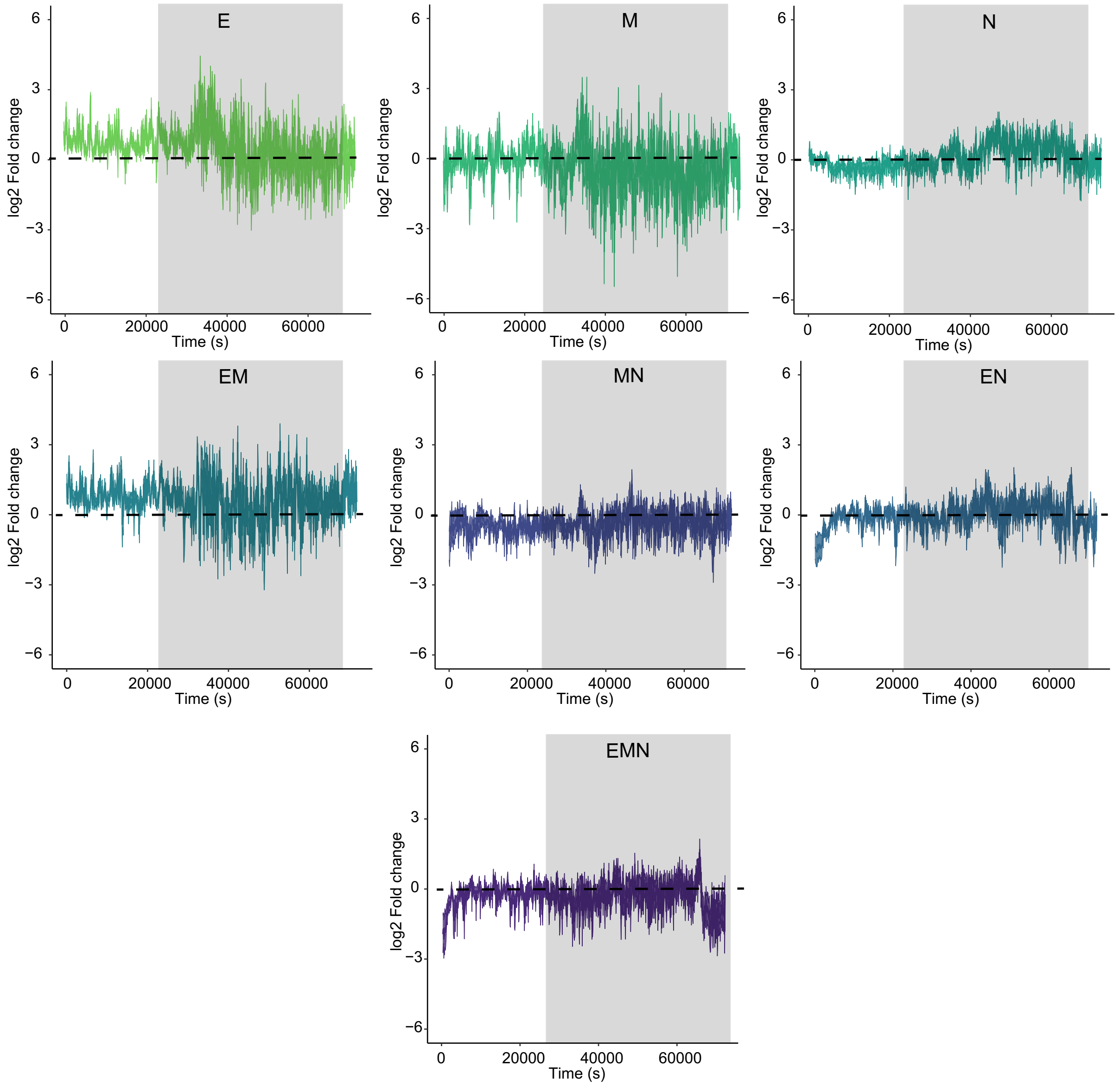


**Supp.Fig.10: Traces of log2 fold change in movement of 14 dpf-larvae throughout the sleep test for each condition relative to controls.**

Average locomotor response of larvae from each group throughout the duration of the sleep test. Larvae exposed to E or EM showed increased activity log2 fold changes compared to controls, especially before the night phase. Sleep-like behavioural patterns were disrupted in the E, M and EM groups, with irregular fluctuations in activity levels observed across the night. n ≥ 24 per condition.

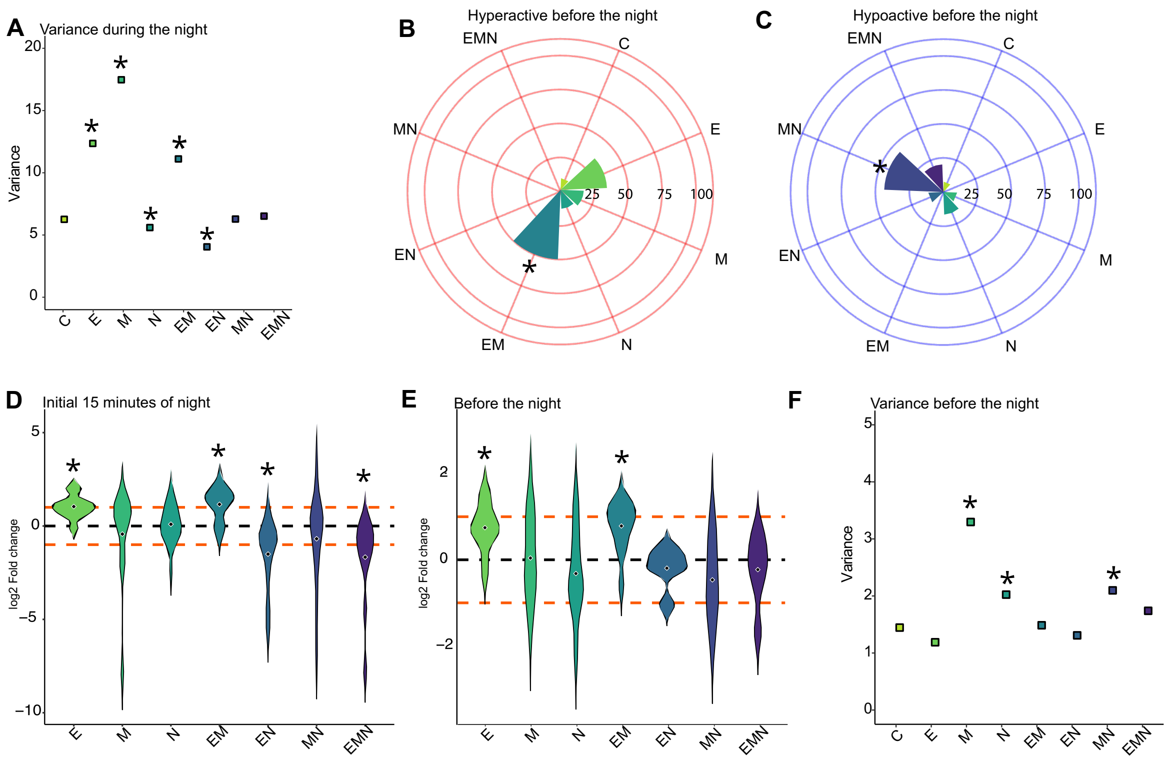


**Supp.Fig.11: Behavioural variability and log2 fold change in movement before and during the night at 14 dpf.**

**A.**Animals from the E, M or EM groups had a greater variance during the night, while those from the N or EN groups had a lower one. * = $p < 0.05$ compared to the control group. Significance was determined using a Levene test followed by a Bonferroni correction for multiple comparisons. **B.** The percentage of larvae with an average log2 fold change of greater than 1 before the dark phase was calculated. * = $p < 0.05$ compared to control animals. P-values were calculated by using a logistic regression. Over 50% of EM-exposed and at least 25% of E-exposed larvae displayed hyperactivity. **C.** The percentage of larvae with an average log2 fold change of lower than -1 before the dark phase was calculated. * = $p < 0.05$ compared to control animals. P-values were calculated by using a logistic regression. MN-exposed animals tended to be hypoactive. **D.** During the initial 15 minutes of the night**,** E and EM-exposed larvae showed heightened activity, while EN- and EMN-exposed animals exhibited reduced movement. **E.** Before the dark phase, E- and EM-exposed animals were more active than controls. **F.** Animals from the M, N, MN and especially EMN had a greater variance before the night.* = $p < 0.05$ compared to the control group. Significance was determined using a Levene test followed by a Bonferroni correction for multiple comparisons. For all experiments, $n \geq 24$ per condition. The violin plot displays the data in log2 fold change relative to control animals. The black dashed line indicates the average of control larvae (0), while the orange lines represent a log2 fold change of +1 and -1. A log2 fold change of +1 represents a twofold increase in movement, while -1 indicates a twofold decrease compared to control animals. This experiment employed a between-subjects design.

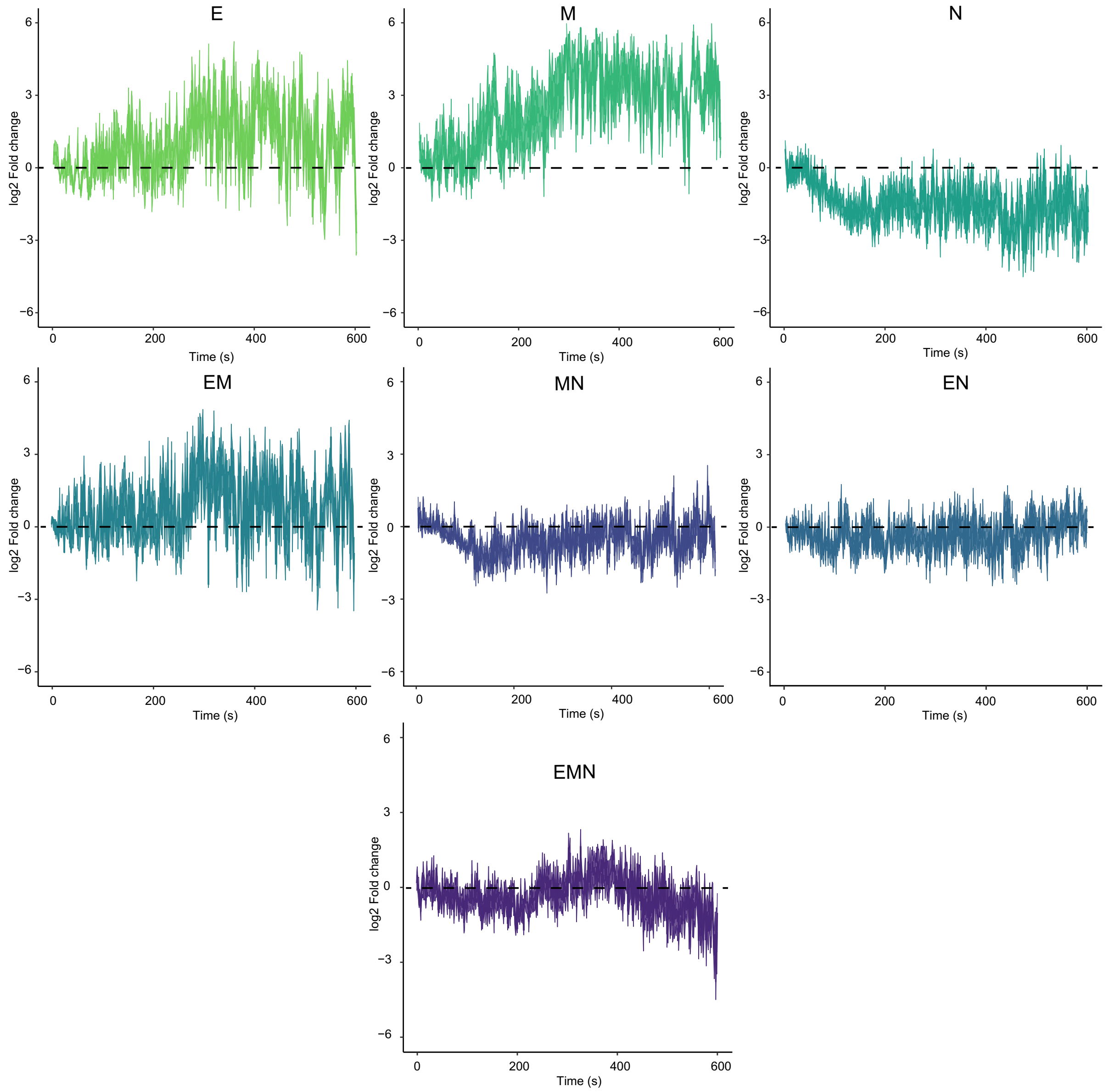


**Supp.Fig.12: Traces of log2 fold change in movement of 8 dpf-larvae across the thermosensation assay for each condition relative to controls.**

Average locomotor response of larvae from each group throughout the duration of the thermosensation test. Larvae exposed to E, M, or their combination showed higher activity log2 fold changes compared to controls, while those exposed to N tended to be hypoactive. Animals in the MN, EN, and EMN groups exhibited responses similar to those of the controls. $n \geq 24$ per condition.

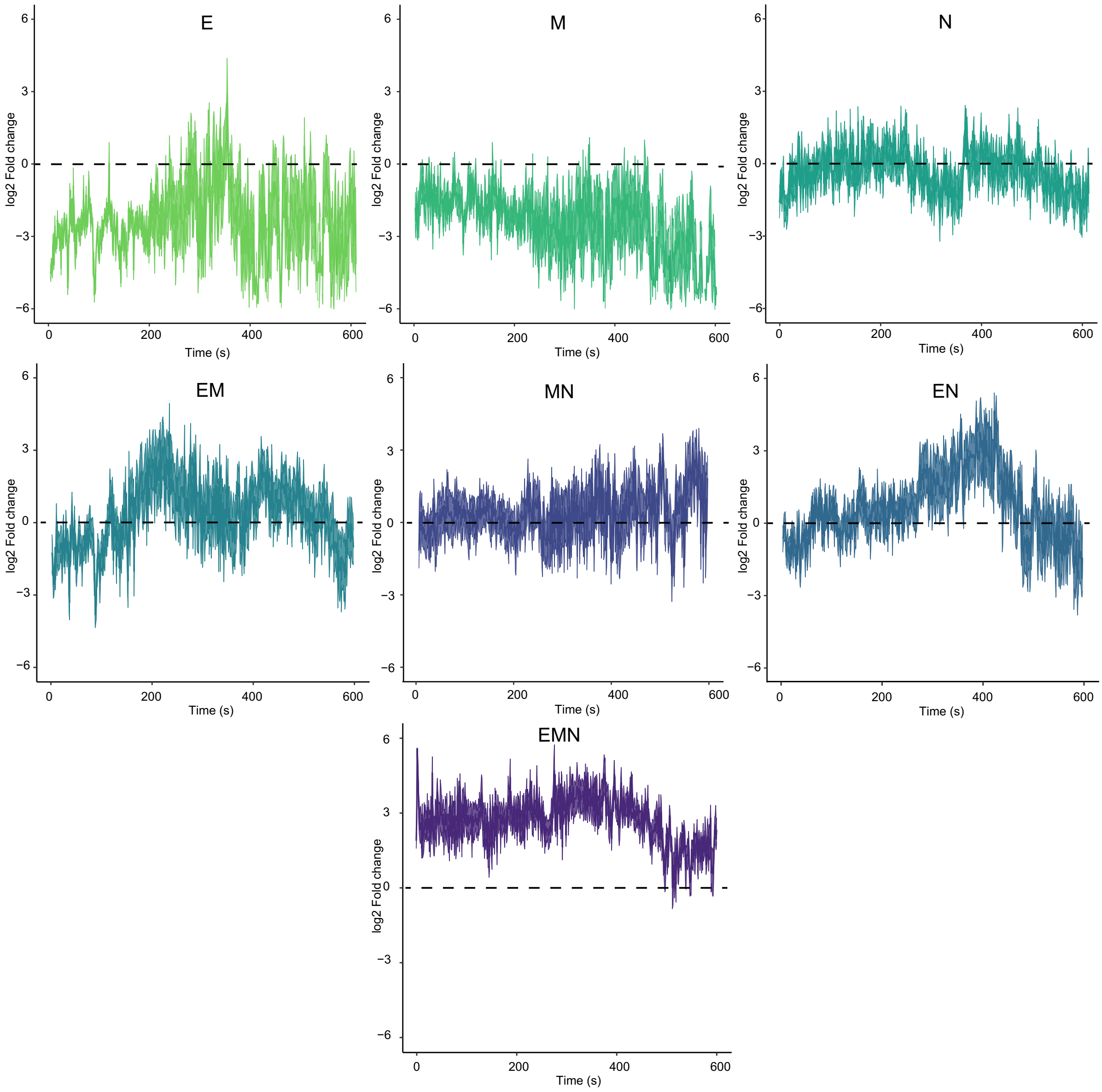


**Supp.Fig.13: Traces of log2 fold change in movement of 14 dpf-larvae across the thermosensation assay for each condition relative to controls.**

Average locomotor response of larvae from each group throughout the duration of the thermosensation test. Larvae exposed to E or M, or their combination, showed decreased activity log2 fold changes compared to controls, while those exposed to EM, EN and EMN tended to be hypoactive. Animals in the MN, EN and EMN groups exhibited higher activity log2 fold changes compared to controls. $n \geq 24$ per condition.

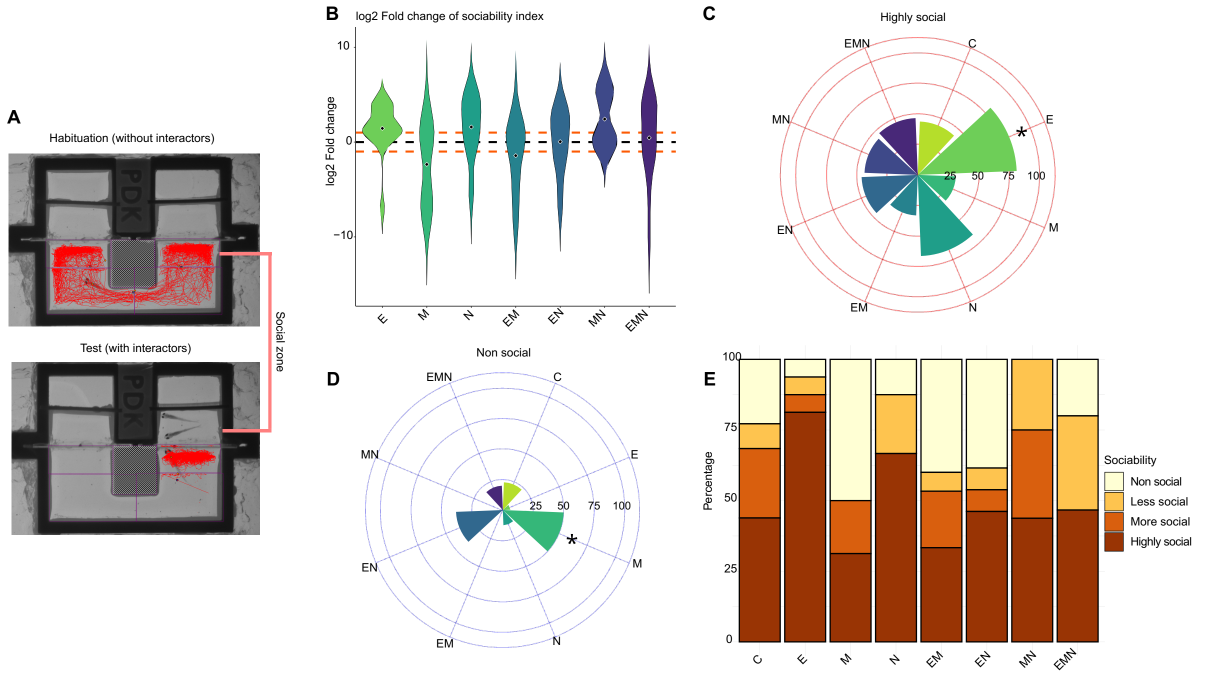


**Supp.Fig.14: Social preference in 21 dpf larvae prenatally exposed**

**A.** Arena used for the social preference test. During the 15-minute habituation session, the tested animal was allowed to explore the arena freely. During the 10-minute test session, two interactors were added to either the top-left or top-right corner. The red trace shows the movement of the tested animal. This example illustrates a highly social individual who spent considerable time interacting with others during the test session. **B.** Larvae exposed to E, N or NM showed increased sociability, while M- and EM- showed decreased sociability compared to controls. No significant differences were observed. **C.** The percentage of larvae with an average log2 fold change of greater than 1 on the social index was calculated. * = $p < 0.05$ compared to control animals. P-values were calculated by using a logistic regression. A large proportion of animals in the E and N groups showed increased sociability. **D.** The percentage of larvae with an average log2 fold change of lower than -1 on the social index was calculated. * = $p < 0.05$ compared to control animals. P-values were calculated by using a logistic regression. The M and EN groups had the highest proportions of individuals with decreased social index. **E.** Manual clustering based on the average log2 fold change on the social index for each larva. The categories were separated as follows: Non-social log2 fold change lower than -1, Less social log2 fold change between -1 and 0, More social log2 fold change between 0 and 1 and Highly social log2 fold change greater than 1. Control animals showed a homogenous distribution. Most E- and N- exposed larvae were highly social. No non-social individuals were found in the MN group. $n \geq 15$ per condition.
